# The global spectrum of tree crown architecture

**DOI:** 10.1101/2024.09.14.613032

**Authors:** Tommaso Jucker, Fabian Jörg Fischer, Jérôme Chave, David A. Coomes, John Caspersen, Arshad Ali, Grace Jopaul Loubota Panzou, Ted R. Feldpausch, Daniel Falster, Vladimir A. Usoltsev, Toby D. Jackson, Stephen Adu-Bredu, Luciana F. Alves, Mohammad Aminpour, Ilondea B. Angoboy, Niels P. R. Anten, Cécile Antin, Yousef Askari, Rodrigo Muñoz, Narayanan Ayyappan, Lindsay Banin, Nicolas Barbier, John J. Battles, Hans Beeckman, Yannick E. Bocko, Ben Bond-Lamberty, Frans Bongers, Samuel Bowers, Michiel van Breugel, Arthur Chantrain, Rajeev Chaudhary, Jingyu Dai, Michele Dalponte, Kangbéni Dimobe, Jean-Christophe Domec, Jean-Louis Doucet, Juan Manuel Dupuy Rada, Remko A. Duursma, Moisés Enríquez, Karin Y. van Ewijk, William Farfán-Rios, Adeline Fayolle, Marco Ferretti, Eric Forni, David I. Forrester, Hammad Gilani, John L. Godlee, Matthias Haeni, Jefferson S. Hall, Jie-Kun He, Andreas Hemp, José L. Hernández-Stefanoni, Steven I. Higgins, Robert J. Holdaway, Kiramat Hussain, Lindsay B. Hutley, Tomoaki Ichie, Yoshiko Iida, Hai-sheng Jiang, Puspa Raj Joshi, Hasan Kaboli, Maryam Kazempour Larsary, Tanaka Kenzo, Brian D. Kloeppel, Takashi Kohyama, Suwash Kunwar, Shem Kuyah, Jakub Kvasnica, Siliang Lin, Emily R. Lines, Hongyan Liu, Craig Lorimer, Jean-Joël Loumeto, Yadvinder Malhi, Peter L. Marshall, Eskil Mattsson, Radim Matula, Jorge A. Meave, Sylvanus Mensah, Xiangcheng Mi, Stéphane T. Momo, Glenn R. Moncrieff, Francisco Mora, Sarath P. Nissanka, Zamah Shari Nur Hajar, Kevin L. O’Hara, Steven Pearce, Raphaël Pelissier, Pablo L. Peri, Pierre Ploton, Lourens Poorter, Mohsen Javanmiri Pour, Hassan Pourbabaei, Sabina C. Ribeiro, Casey Ryan, Anvar Sanaei, Jennifer Sanger, Michael Schlund, Giacomo Sellan, Alexander Shenkin, Bonaventure Sonké, Frank J. Sterck, Martin Svátek, Kentaro Takagi, Anna T. Trugman, Matthew A. Vadeboncoeur, Ahmad Valipour, Mark C. Vanderwel, Alejandra G. Vovides, Peter Waldner, Weiwei Wang, Li-Qiu Wang, Christian Wirth, Murray Woods, Wenhua Xiang, Fabiano de Aquino Ximenes, Yaozhan Xu, Toshihiro Yamada, Miguel A. Zavala, Niklaus E. Zimmermann

## Abstract

Trees can differ enormously in their crown architectural traits, such as the scaling relationships that link their height and crown size to their stem diameter. Yet despite the importance of crown architecture in shaping the structure and function of woody ecosystems, we lack a complete picture of what drives this incredible diversity in crown shapes. Using data from >500,000 globally distributed trees, we explored how climate, disturbance, competition, functional traits, and evolutionary history constrain the height, crown size and shape of the world’s tree species. We found that variation in height scaling relationships was primarily controlled by water availability and light competition. Conversely, crown width was predominantly shaped by exposure to wind and fire, while also covarying with other functional traits related to mechanical stability and photosynthesis. Additionally, several plant lineages had crown architectures that defy their environments, such as the exceedingly slender dipterocarps of Southeast Asia, or the extremely wide crowns of legumes in African savannas. Our study charts the global spectrum of tree crown architectural types. It provides a roadmap for integrating crown architecture with vegetation models and remote sensing observations, so that we may better understand the processes that shape the 3D structure of woody ecosystems.

## INTRODUCTION

Trees come in all shapes and sizes – from incredibly tall and slender, to short with wide, flat crowns ^1–6^. This incredible diversity in tree crown architecture plays an important role in driving variation in growth, water use and competition among tree species ^1,4,7,8^. Moreover, tree crown architecture underpins key emergent properties of woody ecosystems, including their 3D canopy structure, aboveground biomass, primary productivity and hydrology ^7,9–14^. Consequently, uncovering the environmental, ecological and evolutionary drivers that shape the crown architecture of the world’s trees is central to better understanding the processes that constrain the structure and function of woody ecosystems. It is also essential for developing more realistic representations of these ecosystems in vegetation models ^15–18^ and scaling between field and remote sensing observations ^19–22^ – both of which are critical for tracking and predicting how terrestrial ecosystems are responding to rapid global change.

Differences in crown architecture among trees are the result of species employing a variety of strategies to meet a series of competing physiological, structural, competitive, defensive and reproductive demands (Table 1). Trees expand their crowns vertically and laterally to intercept light, compete with neighbours and disperse seeds, while also needing to maintain water transport to their leaves and mechanical stability ^4,6,23–29^. The balance different tree species strike between these various priorities depends on their environment, ecological strategy and evolutionary history, and will be reflected in the scaling relationships between different axes of tree size, such as their height, crown width and stem diameter ^7,8,20,30–38^. For instance, in arid climates woody biomass allocation tends to shift away from height growth to limit the risk of hydraulic failure, resulting in trees that are shorter for a given diameter ^2,30,35,39,40^. Conversely, when water and nutrients are non-limiting, strong competition for light leads to greater investment in height growth, pushing trees closer to their structural and hydraulic safety margins ^7,14,24,25,30^. Similarly, tree species have also adapted the size and shape of their crowns to minimise the risk of damage from wind, fire, snow and browsing ^37,41–46^. Yet despite clear evidence that crown allometric scaling relationships can vary considerably among tree species, we lack a unified picture of how and why they do so. Nor do we understand how different axes of crown size and shape covary with one another, how they relate to other key plant functional traits, or how they vary among plant lineages.

**Table 1:**
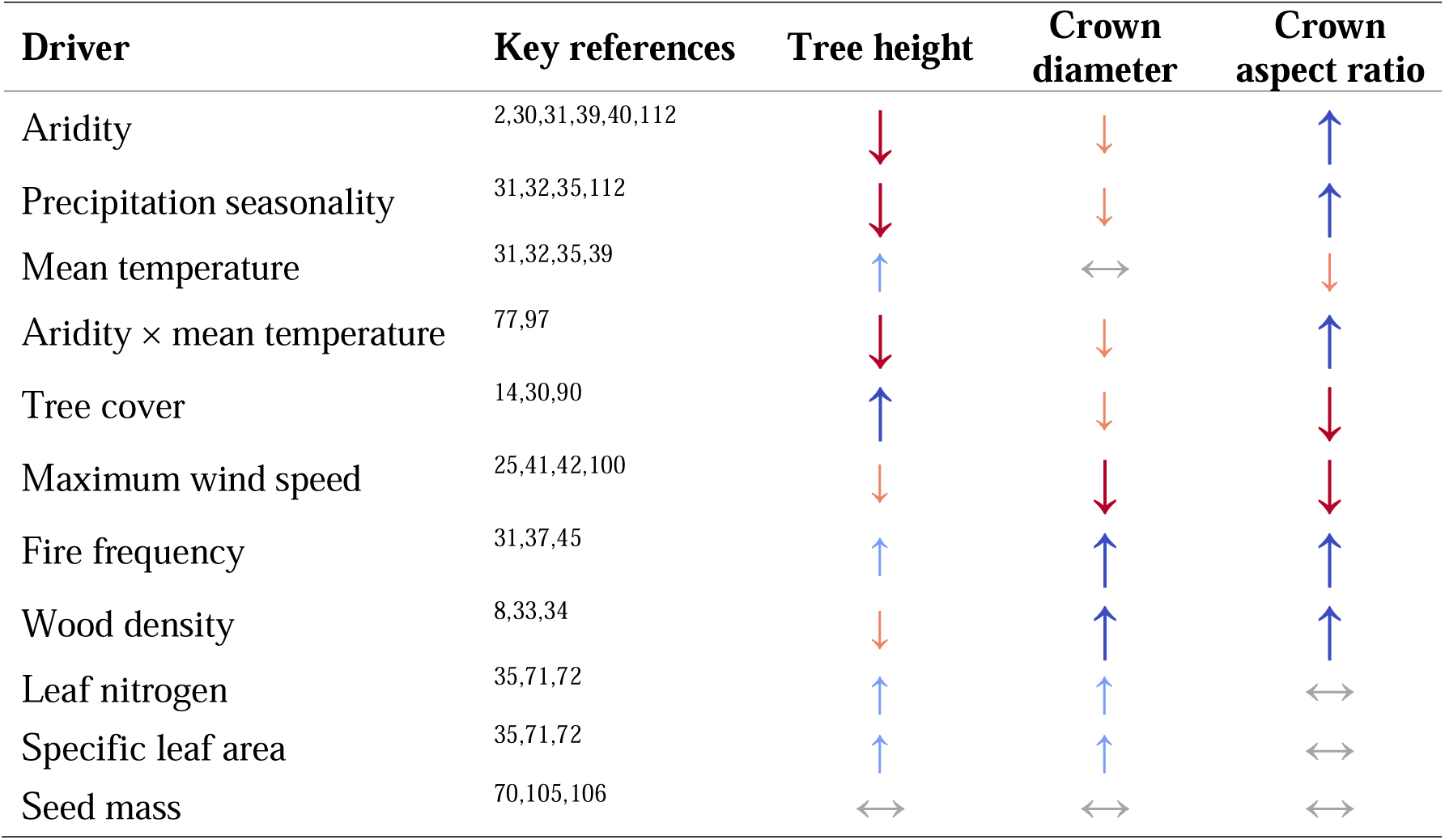
Predicted relationships between size-standardized estimates of tree height, crown diameter and crown aspect ratio (i.e., after controlling for differences in stem diameter) and various climatic drivers, tree cover (as a proxy for competitive environment), disturbance agents, and functional traits. Upward-pointing arrows in blue 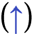 denote positive relationships, while negative ones are shown as downward-pointing red arrows 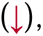 with the size of the arrows reflecting the expected strength of the relationship. Double-headed arrows 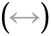 indicate relationships that are expected to be either weak or variable. References supporting each of these hypothesised effects are provided in the table.

| Driver | Key references | Tree height | Crown diameter | Crown aspect ratio |
| --- | --- | --- | --- | --- |
| Aridity | 2,30,31,39,40,112 | ↓ | ↓ | ↑ |
| Precipitation seasonality | 31,32,35,112 | ↓ | ↓ | ↑ |
| Mean temperature | 31,32,35,39 | ↑ | ↔ | ↓ |
| Aridity × mean temperature | 77,97 | ↓ | ↓ | ↑ |
| Tree cover | 14,30,90 | ↑ | ↓ | ↓ |
| Maximum wind speed | 25,41,42,100 | ↓ | ↓ | ↓ |
| Fire frequency | 31,37,45 | ↑ | ↑ | ↑ |
| Wood density | 8,33,34 | ↓ | ↑ | ↑ |
| Leaf nitrogen | 35,71,72 | ↑ | ↑ | ↔ |
| Specific leaf area | 35,71,72 | ↑ | ↑ | ↔ |
| Seed mass | 70,105,106 | ↔ | ↔ | ↔ |

Here we assembled a global dataset capturing information on the stem diameter (*D*), height (*H*), crown diameter (*CD*) and crown aspect ratio (*CAR*, defined as *CD/H*) for over half a million trees (Fig. 1). Using these data, we developed an approach for modelling variation in *H–D*, *CD–D* and *CAR–D* scaling relationships among species that allowed us to compare their crown sizes and shapes while explicitly controlling for differences in their stem sizes. We applied this method to 1914 well-sampled tree species that span all major woody biomes and clades and used it to: (**1**) characterise the full spectrum of crown architectural types observed across the world’s tree species and biomes; (**2**) explore whether crown architectural traits are phylogenetically constrained and identify which clades have crown sizes and shapes that are particularly extreme; and (**3**) test a series of predictions about how *H–D*, *CD–D* and *CAR–D* scaling relationships vary in relation to climate, competition, disturbance and other functional traits related to plant metabolism, hydraulics, structural stability and dispersal (Table 1).

**Fig. 1:**
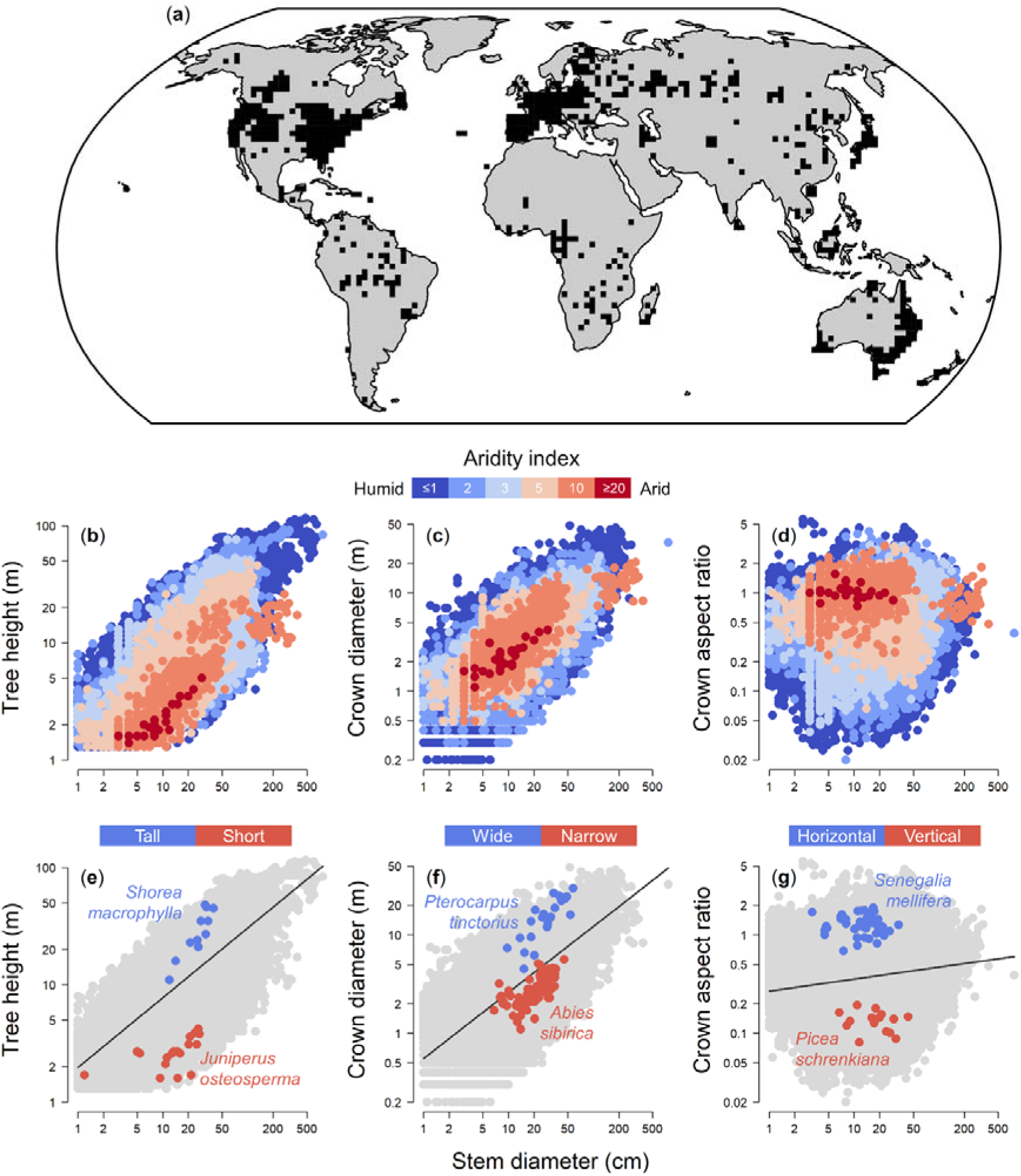
Overview of the global tree allometry dataset. A world map shows the geographic distribution of the allometric data (**a**), with locations from which individual tree records were sourced shown in black (data aggregated in 200×200 km grid cells). The relationship between each tree’s stem diameter and its (**b**) height (*H*), (**c**) crown diameter (*CD*) and (**d**) crown aspect ratio (*CAR*) is shown on a logarithmic scale (*n* = 374,888 individual trees belonging to 1914 species). *CAR* is defined as the ratio between *CD* and *H*, with values lower than one indicating a vertical crown profile (*H* > *CD*) while values greater than 1 corresponding to a horizontal crown profile (*CD* > *H*). Points in (**b–d**) are coloured according to the aridity index value assigned to each tree based its geographic coordinates, with larger values corresponding to drier conditions (shown in red). Panels (**e–g**) illustrate the approach used to generate size-standardized estimates of *H*, *CD* and *CAR* for each tree species. Black lines are predicted values obtained by fitting a linear regression to the entire dataset (grey points). By comparing predicted and observed value of *H*, *CD* and *CAR*, we quantified how much each species departs on average from this general trend and identified ones with greater (blue points) or smaller *H*, *CD* and *CAR* values (red points) than expected given their stem diameters.

## MATERIAL AND METHODS

### Individual tree height and crown size data

We compiled 528,311 georeferenced records of individual trees for which stem diameter (*D*, cm), height (*H*, m) and/or crown diameter (*CD*, m) were measured (Fig. 1a). For trees where both *H* and *CD* were measured (*n* = 340,221; 64.4%), we also calculated their crown aspect ratio (*CAR*) as *CD*/*H*, where *CAR* < 1 denotes a vertical crown profile and *CAR* > 1 a flat or horizontal profile ^1,47^. These data were obtained from 62,435 globally distributed sites which encompass all major terrestrial biomes and span a gradient in mean annual temperature of - 15.1–30.1°C and 143–7157 mm yr^-1^ in rainfall. Sampled trees span multiple orders of magnitude in size and crown shape (Fig. 1b–d) and represent 5161 tree species from 1451 genera and 187 plant families.

Most of the data (94.4% of records) were sourced from the Tallo database ^2^. Additionally, we also obtained data from Alberta’s Permanent Sample Plots network in Canada (*n* = 12,171 trees) and the ICP Forests network in Europe (*n* = 17,540 trees). Allometric data were quality controlled following the protocols of the Tallo database ^2^. Briefly, we first used Mahalanobis distance to identify and remove possible data entry errors by screening for trees with unrealistically large or small *H* and *CD* values for a given stem diameter. Species names were then standardized against those of The Plant List (TPL) using the *taxonstand* package ^48^ in R (version 4.1.1) ^49^. Lastly, we excluded records from species that did not meet our working definition of trees: perennial woody seed plants with a single dominant stem that are self-supporting and undergo secondary growth (i.e., excluding ferns, palms, short multi-stemmed shrubs and lianas).

### Species level, size-standardized estimates of tree height, crown diameter and crown aspect ratio

Tree species that can differ considerably in their maximum size and developmental strategies, so to directly compare their crown architecture we used two complimentary approaches to generate size-standardized estimates of *H*, *CD* and *CAR* at the species level. For these and all subsequent analyses, we focused on species with at least 10 trees sampled within the same biome and spanning a minimum *D* range of 20 cm between the smallest and largest measured tree (see ‘*Environmental data*’ section below for details on how trees were assigned to biomes). In total, 1914 species represented by 374,888 individual trees met these criteria for at least one of the three axes of crown size and shape and 1309 species represented by 251,733 trees had sufficient data for all three (1910 species for *H*, 1313 for *CD* and 1309 for *CAR*; see Table S1 in Supporting Information for details). These 1914 species cover 755 genera and 131 plant families.

The first approach to comparing species’ crown architectures involved generating estimates of *H*, *CD* and *CAR* for a tree of fixed size (*D* = 30 cm) for each species ^30,50^. To do this, we modelled variation in *H*, *CD* and *CAR* among individual trees as a power-law function of *D* by fitting linear mixed-effects regressions to log-log transformed data ^7,30,31^:

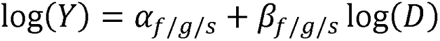

where *Y* denotes either *H*, *CD* or *CAR*, α is the intercept (or normalization constant) and β is the slope (or scaling exponent). Models were fit using the *lme4* package ^51^ and both the intercept and slope of the regressions were allowed to vary among tree species, genera and families with a nested random effects structure (denoted by the *f/g/s* subscripts in the equation above). *R^2^* values accounting for both fixed and random effects components of the models were 0.82, 0.74 and 0.45 for *H*, *CD* and *CAR*, respectively ^52^. The fitted models were then used to predict *H*, *CD* and *CAR* for each species assuming a fixed stem size of 30 cm (hereafter *H_D=30_*, *CD_D=30_* and *CAR_D=30_*).

The second approach we developed to capture variation in crown architecture among species is conceptually similar to the first, but avoids the need to choose an arbitrary stem size at which to compare species ^7^. As for the previous method, we began by using the tree-level data to model variation in *H*, *CD* and *CAR* as a power-law function of *D*. However, in this case we explicitly ignored differences among species and simply estimated the overall scaling relationships between *H*–*D*, *CD*–*D* and *CAR*–*D* across the whole dataset by fitting ordinary linear regressions to log-log transformed data. Using the residuals of the models (i.e., the difference between observed and predicted values of *H*, *CD* and *CAR* on a log-log scale), we then determined *post hoc* whether a given species has *H*, *CD* and *CAR* values that are – on average – larger (positive residuals) or smaller (negative residuals) than expected after accounting for differences in stem size between trees (see Fig. 1e–g for a graphical representation of this approach). Because sample sizes varied considerably among species (*n* = 10–22,835 trees per species), we subset the data by randomly selecting 10 trees per species prior to model fitting. Without this step, well-sampled species would dominate the signal of the regression and skew the values of the residuals. This randomization step was repeated 100 times and for each species we then calculated the mean value of the residuals across all model runs as a measure of size-standardized *H*, *CD* and *CAR* (hereafter *H_RESID_*, *CD_RESID_* and *CAR_RESID_*).

Quantitatively, the two approaches gave very similar results (Fig. S1). However, the second method based on model residuals is better suited to comparing species that exhibit contrasting growth trajectories or vary in their size at maturity, as it integrates data across all observed tree sizes instead of focusing on a single point of comparison (e.g., *D* = 30 cm, which could correspond to a small tree for some species and a very large one for others). For subsequent analyses we therefore focus on comparing values of *H_RESID_*, *CD_RESID_*and *CAR_RESID_* across species, but to aid the interpretation of results we also report values of *H_D=30_*, *CD_D=30_*and *CAR_D=30_*.

### Environmental data

To understand how environmental conditions shape variation in *H_RESID_*, *CD_RESID_* and *CAR_RESID_* among tree species, we used the geographic coordinates of individual trees to assign attributes related to climate, competition and disturbance (see Table S2 for full details on sources of environmental data). These environmental predictors were chosen based on previous work suggesting they play an important role in shaping tree crown allometry by constraining plant hydraulics, growth and competition (Table 1). Importantly, they were also selected as they were not strongly correlated with one another (Fig. S2), allowing their effects to be teased apart in subsequent analyses. In addition to the environmental predictors described below, trees were also assigned to one of seven biome classes based on the classification used by the Terrestrial Ecoregions of the World database ^53^.

For climate we focused on the effects of mean annual temperature (MAT, °C), precipitation seasonality (mm) and aridity (unitless index). MAT and precipitation seasonality were obtained from the WorldClim2 database at a resolution of 30 arc-seconds ^54^. Aridity was instead calculated as the ratio between potential evapotranspiration (PET, mm) and mean annual precipitation (MAP, mm), where MAP was obtained from WorldClim2 while PET was derived from the Global Aridity Index and Potential Evapotranspiration Climate Database at 30 arc-second resolution ^55^. This is the inverse of how aridity is often expressed ^55^, but has the advantage of being easier to interpret as larger values of the aridity index correspond to drier conditions ^56^.

As a proxy for local competitive environment, we used estimates of tree cover derived from MODIS at 15-arc second resolution for the year 2008 ^57^. We chose this approach as for most trees we lacked information on stand-level attributes commonly used to characterise competition, such as basal area or stem density ^14,30^. However, for a subset of sites across which these field data were available, we found good agreement between MODIS-derived estimates of tree cover and stand basal area (Fig. S3), suggesting satellite estimates of tree cover provide a reliable indicator of competitive environment.

As indicators of disturbance, we focused on wind speed and fire risk. Specifically, we used the ERA5-Land data to calculate the maximum wind gust speed experienced by each tree between 2010–2020. To quantify exposure to fire we calculated the mean burned area fraction between 2001–2010, as estimated from MODIS in the Global Fire Emissions Database ^58^. Note that we also tried to assess the impacts of snow accumulation on tree crowns, but found that estimates of snow cover duration derived from MODIS were strongly correlated with MAT (Pearson correlation coefficient, ρ = -0.80; Fig. S2), and where therefore not considered in subsequent analyses.

For each of the 1914 species that met the minimum sampling criteria described previously, we then calculated mean values of aridity, MAT, precipitation seasonality, tree cover, maximum wind gust speed and burned area fraction across all sampled trees. Each species was also assigned to a unique biome based on the terrestrial ecoregion in which they were recorded most frequently ^59^.

### Functional trait data

To test how variation in *H_RESID_*, *CD_RESID_* and *CAR_RESID_* among tree species relates to other key plant functional traits, we compiled data on wood density (g cm^-3^), leaf nitrogen content (mg g^-1^), specific leaf area (SLA, mm^2^ mg^-1^) and seed mass (g) from multiple sources (Table S3). This includes the TRY plant trait database ^60^, the Botanical Information and Ecology Network (BIEN) database ^61^, the global wood density database ^62^, the Royal Botanic Gardens Kew seed information database, the AusTraits database ^63^, the China plant trait database ^64^, the Terrestrial Ecosystem Research Network (TERN), as well as selected publications ^33,65–69^. These four functional traits were chosen as previous work suggests they may covary with crown size and shape through their influence on whole-plant growth, size, hydraulics and mechanical stability ^30,33,34,70–72^ and have been measured for numerous tree species.

To obtain species-level mean values for each trait, we first grouped together individual records by site (based on shared geographic coordinates) and then species ^73^. For species where no individual-level records could be sourced, species-level values reported in the literature were used instead if available. Of the 1914 tree species for which we estimated *H_RESID_*, *CD_RESID_* and/or *CAR_RESID_*, we obtained wood density estimates for 1572 species (82%), leaf nitrogen content for 1085 (57%), SLA for 1120 (59%) and seed mass for 1108 (58%).

### Mapping the spectrum of crown architectural types and its distribution across woody biomes

To characterise the range of crown forms that tree species can assume and better understand how these vary among woody biomes, we used estimates of *H_RESID_*, *CD_RESID_* and *CAR_RESID_* to determine how species cluster into architectural types based on their height, crown size and shape ^1^. To do this, we first calculated the correlation between *H_RESID_* and *CD_RESID_* to determine how tightly constrained these two axes of crown architecture are across the 1309 species where both had been measured. A strong positive correlation would indicate that species which are taller for a given stem diameter also tend to have larger crowns and *vice versa*. Conversely, a weak correlation between *H_RESID_* and *CD_RESID_* would suggest that when standardized by size, tree species are able to adopt a wide range of crown architectural forms, from tall and narrow to short and wide.

We then grouped species into one of nine crown architectural types: (1) short and narrow, (2) narrow, (3) tall and narrow, (4) short, (5) medium-sized, (6) tall, (7) short and wide, (8) wide, and (9) tall and wide species. Species were assigned to groups based on their *H_RESID_*and *CD_RESID_* values and whether these fell in the lower quartile, interquartile range or upper quartile of data (see Fig. 2a for a visual representation). Note that *CAR_RESID_*was not used to group species, as any differences in *CAR_RESID_* among species can be directly attributed to ones in *H_RESID_* and *CD_RESID_*. To determine the degree to which crown forms are adapted and confined to specific environments, we quantified the relative frequency of each architectural type across different biomes. To support this analysis, we also used one-way ANOVAs to compare mean values of *H_RESID_*, *CD_RESID_* and *CAR_RESID_* among species from different biomes.

**Fig. 2:**
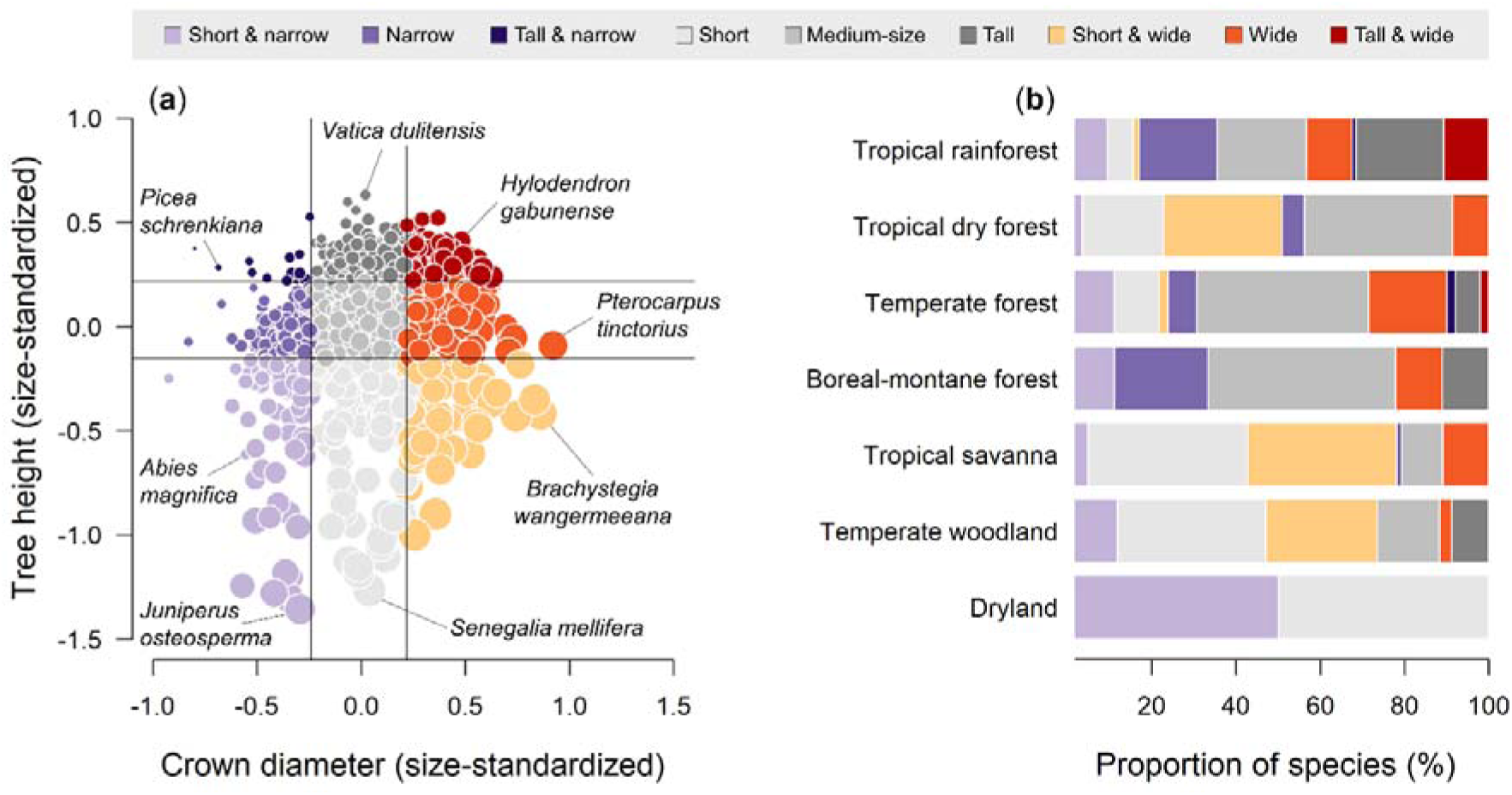
Tree crown architectural types (**a**) and their distribution across biomes (**b**) for the 1309 tree species for which both height and crown size were measured. Tree species were grouped into one of nine architectural types based on their size-standardized height (*H_RESID_*) and crown diameter values (*CD_RESID_*). In (**a**) the vertical and horizontal lines mark the 25^th^ and 75^th^ percentile of the data and the size of each circle reflects the crown aspect ratio (*CAR_RESID_*). Examples of tree species that occupy different areas of this crown architectural spectrum are highlighted. In (**b**) the proportion of species belonging to the nine architectural types is reported for each biome (see Table S6 for pairwise comparisons of *H_RESID_*, *CD_RESID_* and *CAR_RESID_*values among biomes, and Fig. S4 for a breakdown of the nine architectural types among angiosperms and gymnosperms).

### Evolutionary history and its fingerprint on crown architecture

To determine whether crown architectural traits exhibit phylogenetical signal, we mapped *H_RESID_*, *CD_RESID_* and *CAR_RESID_* onto the Smith & Brown (2018) phylogeny of seed plants and calculated Pagel’s λ as a general test of phylogenetic signal for each crown attribute ^74,75^. A λ value of 0 indicates no phylogenetic signal while a value of 1 corresponds to a trait that has evolved according to Brownian motion, indicating strong phylogenetic signal ^74^. Pagel’s λ was calculated using the *phytools* package ^76^, which uses a likelihood ratio test to determine whether λ is significantly different from 0. Because tests of phylogenetic signal are sensitive to errors in the phylogeny, such as those associated with branch lengths ^75^, only species that were a direct match to those in the time-calibrated phylogeny were retained for subsequent analyses (1225 species for *H_RESID_*, 870 for *CD_RESID_* and 868 for *CAR_RESID_*; Table S1).

To complement λ – which provides a global test of phylogenetic signal across the entire phylogeny – we also explored how *H_RESID_*, *CD_RESID_* and *CAR_RESID_* varied among clades within the phylogeny. Specifically, we used one-way ANOVAs fit without an intercept to identify plant families and genera where species’ mean *H_RESID_*, *CD_RESID_* and *CAR_RESID_* values are significantly greater or smaller than zero. For this purpose, we only retained families and genera represented by at least five species in our dataset (*n* = 63 families and 85 genera for *H_RESID_*, and *n* = 56 families and 60 genera for *CD_RESID_* and *CAR_RESID_*).

### Effects of climate, competition, disturbance and functional traits on crown architecture

To quantify the effects of climate, competition and disturbance on tree crown architecture, we modelled variation in *H_RESID_*, *CD_RESID_* and *CAR_RESID_*among species as a function of aridity, MAT, precipitation seasonality, tree cover, maximum wind gust speed and burned area fraction using multiple regression. Models also included an interaction term between aridity and MAT to test whether the effects of low water availability on tree height, crown size and shape would be strongest in hotter environments ^77^, as well as a binary variable testing for systematic differences in *H_RESID_*, *CD_RESID_*and *CAR_RESID_* between angiosperms and gymnosperms ^8,30,35^.

To determine whether *H_RESID_*, *CD_RESID_*and *CAR_RESID_* also vary in relation to species’ functional traits, we then fit separate models in which either wood density, leaf nitrogen content, SLA or seed mass were added to the multiple regression alongside the environmental predictors described above. Note that models including functional traits as predictors were restricted to the subset of species for which trait data were available (see Table S1 for details).

Prior to model fitting, both aridity and seed mass were log-transformed to linearise relationships between response and predictor variables. All continuous predictor variables were then centred and scaled by subtracting the mean and dividing by 1 standard deviation, while the binary variable grouping species into major evolutionary clades was coded as –1 for gymnosperms and 1 for angiosperms. This allowed us to directly compare the effect sizes of different predictors both within and across models based on their regression coefficients. To ensure model coefficients were not affected by collinearity among predictors, we calculated variance inflation factors for all models to confirm they were all ≤ 2.

To account for non-independence among species due to shared evolutionary history, regression models were fit using phylogenetic generalised least squares (PGLS) ^78^. PGLS models were fit using the *gls* function in the *nlme* package ^79^, where the correlation structure among species was captured using Pagel’s λ as implemented by the *corPagel* function in the *ape* package ^80^. A phylogenetic tree capturing evolutionary relationships among species was generated using the *V.PhyloMaker* package ^81^, which uses a comprehensive time-calibrated phylogeny of 79,881 seed plant species as a backbone ^82^. Model *R^2^* values that account for the phylogenetic structure of the data were calculated using the *rr2* package ^83^.

## RESULTS

### Global variation in crown architectural types

Across the 1914 tree species considered in our analysis, we found enormous variation in size-standardized estimates of tree height, crown diameter and crown aspect ratio, both when considering values of *H_RESID_*, *CD_RESID_* and *CAR_RESID_* (Fig. 2) and *H_D=30_*, *CD_D=30_* and *CAR_D=30_*. Specifically, *H_D=30_* varied 12.1-fold across species, ranging from < 4 m in species like *Juniperus osteosperma* and *Maerua crassifolia* to > 30 m in several species of the genera *Shorea*, *Parashorea*, *Hopea* and *Vatica* (all Dipterocarpaceae) and as much as 43.2 m in *Eucalyptus regnans*. By contrast, *CD_D=30_* was less than half as variable among species, ranging 5.4-fold from < 4 m in several species of the genus *Picea* to > 14 m in ones like *Brachystegia wangermeeana* and *Pterocarpus tinctorius* in the Fabaceae. As for crown profile shape, *CAR_D=30_*ranged 10.5-fold across species. At one end of the spectrum, species like *Abies sibirica* and *Vatica dulitensis* were more than five times as tall as their crowns are wide (*CAR_D=30_* < 0.2), whilst several species of the genera *Acacia*, *Vachellia* and *Senegalia* (all Fabaceae) had crowns that are noticeably wider than they are tall (*CAR_D=30_* > 1.2).

For the 1309 species for which we were able to estimate both *H_RESID_* and *CD_RESID_*, we found that these two axes of crown architecture were positively correlated with one another (ρ = 0.26, *P* < 0.001). However, despite a general trend of taller trees also having wider crowns, the relationship between *H_RESID_* and *CD_RESID_* was relatively weak. Species occupied all possible combinations of the tree height *vs* crown diameter spectrum (Fig. 2a), including short and narrow (9.3% of species), short and wide (5.4%), tall and wide (7.9%), and tall and narrow (1.0%).

Where species fell within this spectrum depended, at least in part, on their biome association (Fig. 2b). For example, a high proportion of species found in drylands, temperate woodlands, tropical savannas and tropical dry forests had either short and/or wide crowns (59.6–100% of species, depending on the biome), but almost none were tall (0–8.8%). By contrast, boreal, temperate and tropical rainforests had a considerably higher proportion of species with tall and/or narrow crowns (28.0–60.5%). Overall, differences among biomes were more pronounced for height and crown aspect ratio than for crown width, with biome association explaining 33%, 39% and 5% of the variation in *H_RESID_*, *CAR_RESID_* and *CD_RESID_* among species, respectively (see Table S4 for pairwise comparisons among biomes based on ANOVAs). However, our results also revealed considerable variation in tree architectural types within biomes (Fig. 2b), highlighting how tree species with very different crown architectures can be found in very similar environments.

### The fingerprint of evolutionary history and biogeography on crown architecture

From a macroevolutionary perspective, angiosperms and gymnosperms had very similar mean values of *H_D=30_* (17.8 m and 17.3 m, respectively). However, angiosperms spanned a considerably larger range of heights (3.9–42.3 m), particularly at the tall end of the spectrum where they included 99 of the 100 species with the highest *H_D=30_* values. By contrast, both crown diameter and aspect ratio were noticeably larger in angiosperms compared to gymnosperms (*CD_D=30_* = 6.4 m *vs* 5.3 m; *CAR_D=30_* = 0.43 *vs* 0.36). When we placed estimates of *H_RESID_*, *CD_RESID_* and *CAR_RESID_*onto a time-calibrated phylogeny of seed plants we found that all three exhibited a significant degree of phylogenetic signal (Fig. 3), with Pagel’s λ values of 0.70, 0.54 and 0.63, respectively (*P* < 0.001 in all cases).

**Fig. 3:**
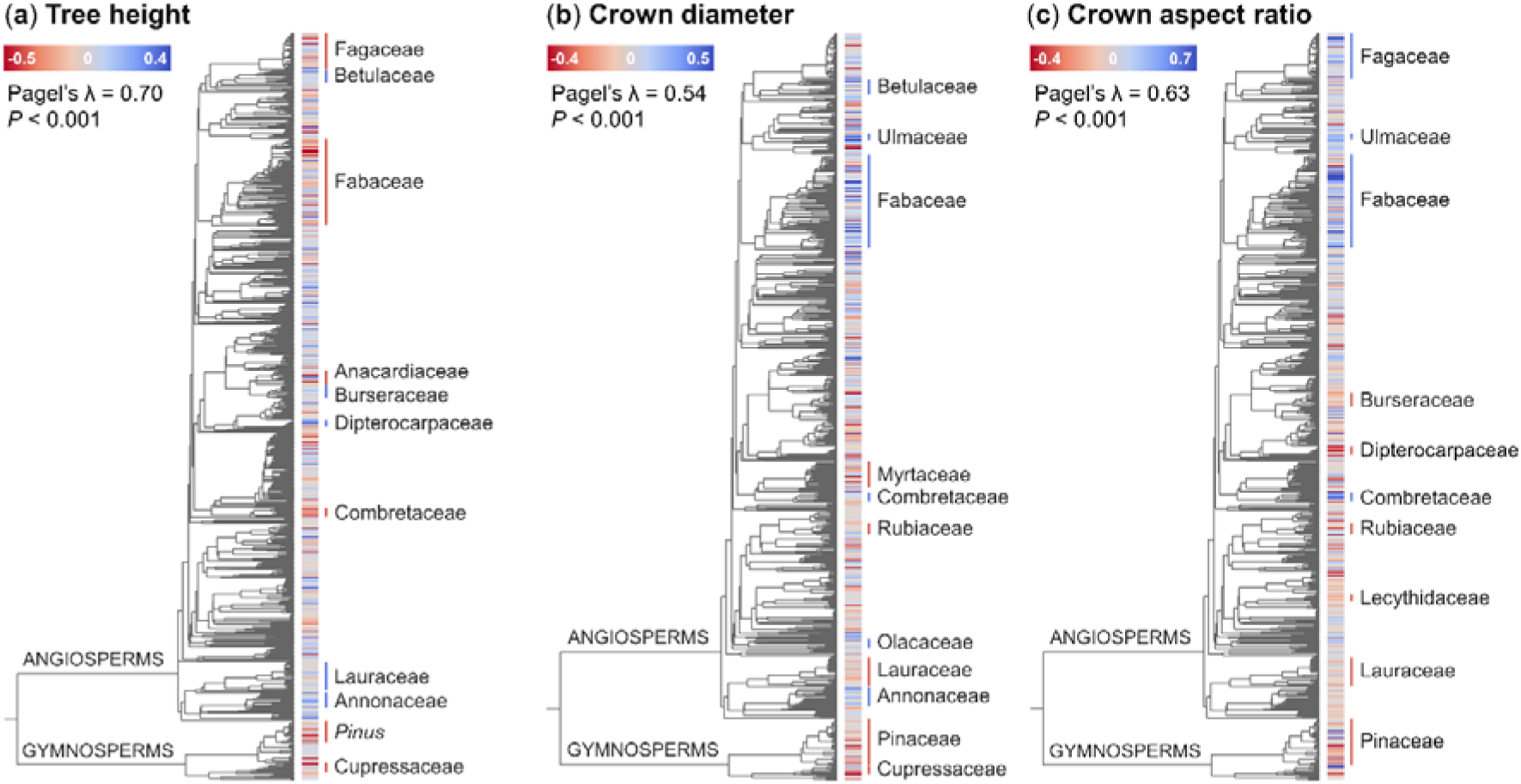
Variation in tree crown architecture across the tree of life. Size-standardized estimates of (**a**) tree height (*H_RESID_*, *n* = 1225 species), (**b**) crown diameter (*CD_RESID_*, *n* = 870 species) and (**c**) crown aspect ratio (*CAR_RESID_*, *n* = 868 species) are mapped onto a time-calibrated phylogeny of seed plants ^82^. Low values (red) indicate species that are shorter, with narrower crowns and smaller crown aspect ratios than expected given the size of their stem, while high values (blue) indicate the opposite. *H_RESID_*, *CD_RESID_*and *CAR_RESID_* all exhibited phylogenetic signal, with Pagel’s λ values of 0.70, 0.54 and 0.63, respectively (*P* < 0.001 in all three cases based on a likelihood ratio test). Plant families and genera with mean *H_RESID_*, *CD_RESID_*and *CAR_RESID_* values that are significantly lower (red lines) or higher (blue lines) than zero are highlighted on each phylogenetic tree (see Tables S4–5 for full details). Note that only species that were a direct match to those in the phylogeny were used for the phylogenetic analysis.

In particular, we found several plant genera and families that stand out based on their tree height, crown size and shape (Fig. 3 and Tables S5–6). For tree height, 25 out of 63 families and 31 out of 86 genera that we tested had mean *H_RESID_* values that were significantly different from zero. Within the angiosperms, species in the Dipterocarpaceae (*H_D=30_* = 24.1 m), Myristicaceae (*H_D=30_* = 24.4 m), Burseraceae (*H_D=30_*= 21.3 m), Annonaceae (*H_D=30_* = 21.3 m) and Betulaceae (*H_D=30_* = 19.6 m) were particularly tall for their stem diameters (Fig. 3a). From a biogeographic standpoint, we found that Southeast Asia was home to an especially high concentration of species with tall and slender growth forms, with nine of the 10 species with the highest *H_RESID_* values native to this region, including species in the genera *Shorea*, *Parashorea* and *Vatica* (Dipterocarpaceae) and *Knema* (Myristicaceae). At the opposite end of the spectrum, species in the Ericaceae (*H_D=30_* = 11.4 m), Combretaceae (*H_D=30_* = 13.3 m), Fagaceae (*H_D=30_* = 15.1 m) and Fabaceae (*H_D=30_* = 16.2 m) were significantly shorter that average for a given stem diameter.

The picture within gymnosperms was equally varied. Cupressaceae were generally shorter than expected (*H_D=30_* = 14.8 m), despite including species like *Sequoia sempervirens* which can grow incredibly tall in absolute terms. Conversely, within the Pinaceae we found a clear divide between species in the genus *Pinus* which are shorter than average (*H_D=30_* = 16.2 m) and those belonging to *Larix* and *Picea* that are taller (*H_D=30_* = 22.2 m and 20.6 m, respectively).

In terms of crown size and shape, we found that 21/56 families and 26/60 genera (*CD_RESID_*) and 12/56 families and 16/60 genera (*CAR_RESID_*) had values that departed significantly from zero (Tables S5–6). One clade that stood out in particular is the Fabaceae (Fig. 3b–c). Of the top 10 species with the highest *CD_RESID_* and *CAR_RESID_*values, four and six were Fabaceae, respectively. Fabaceae had crowns that are both much wider than average (*CD_D=30_* = 7.6 m) and more horizontal in their aspect ratio (*CAR_D=30_* = 0.58). This trend was predominantly driven by species that occupy savannas in Africa and the Americas, including ones in the genera *Senegalia*, *Acacia*, *Brachystegia*, *Vachellia* and *Pterocarpus* (mean *CD_D=30_*and *CAR_D=30_* across 40 tropical savanna specialists = 8.6 m and 0.88, respectively). By contrast, Fabaceae from tropical rainforests (*CD_D=30_* = 7.2 m; *CD_D=30_*= 0.39) and temperate forests (*CD_D=30_* = 6.5 m; *CD_D=30_* = 0.37) had crown sizes and profiles that were very similar to other species in these biomes.

Within gymnosperms, species in the Podocarpaceae (*CD_D=30_* = 5.2 m), Cupressaceae (*CD_D=30_* = 5.3 m) and Pinaceae (*CD_D=30_* = 5.3 m) all exhibited narrower than average crowns (Fig. 3b). In the Pinaceae this effect was far stronger than any variation observed in tree height, resulting in crown aspect ratios that are also much smaller than average (*CAR_D=30_* = 0.34; Fig. 3c). A similar trend emerged for the Rubiaceae and Lauraceae, where species were generally both tall and with narrow crowns, resulting in low crown aspect ratios (*CAR_D=30_* = 0.35 and 0.37, respectively). The exact opposite was true for species in the Combretaceae, Ulmaceae and Fagaceae, which due to their relatively short stature and wide crowns had particularly large crown aspect ratios (*CAR_D=30_* = 0.73, 0.55 and 0.48, respectively).

### Effects of climate, competition, disturbance and functional traits on crown architecture

PGLS models relating variation in tree crown architecture among species to climate, tree cover, disturbance and functional traits explained 55%, 28% and 34% of the variation in *H_RESID_*, *CD_RESID_*and *CAR_RESID_*, respectively (Fig. 4). Differences in height among species were predominantly controlled by water availability (aridity, and to a lesser extent rainfall seasonality) and tree cover (Fig. 4a), with *H_RESID_* decreasing rapidly with rising aridity and increasing steadily as tree cover increased (Fig. 5a–b). On average, species growing where PET was equal to or less than MAP (aridity index ≤ 1) where almost twice as tall for a given stem diameter (*H_D=30_* = 20.1 m) as those where the aridity index was ≥ 2 (*H_D=30_*= 11.1 m). Similarly, *H_D=30_* increased from 11.5 m to 21.4 m when comparing species growing where tree cover was ≤ 20% and ≥ 80% (Fig. 5a). However, we also found that when water was non-limiting (aridity index ≤ 1), trees could vary hugely in their investment in height growth, with *H_D=30_* ranging anywhere between 6.4 m and 43.2 m (Fig. 5b). In contrast to aridity and tree cover, we only observed a modest positive association between *H_RESID_*and MAT (Fig. 4a). MAT did however indirectly influence *H_RESID_* through its interaction with aridity (Fig. S5). Specifically, we found that *H_RESID_*declines with increasing aridity were much less pronounced for trees growing in cold climates (MAT < 10°C) compared to warm ones (MAT > 20°C).

**Fig. 4:**
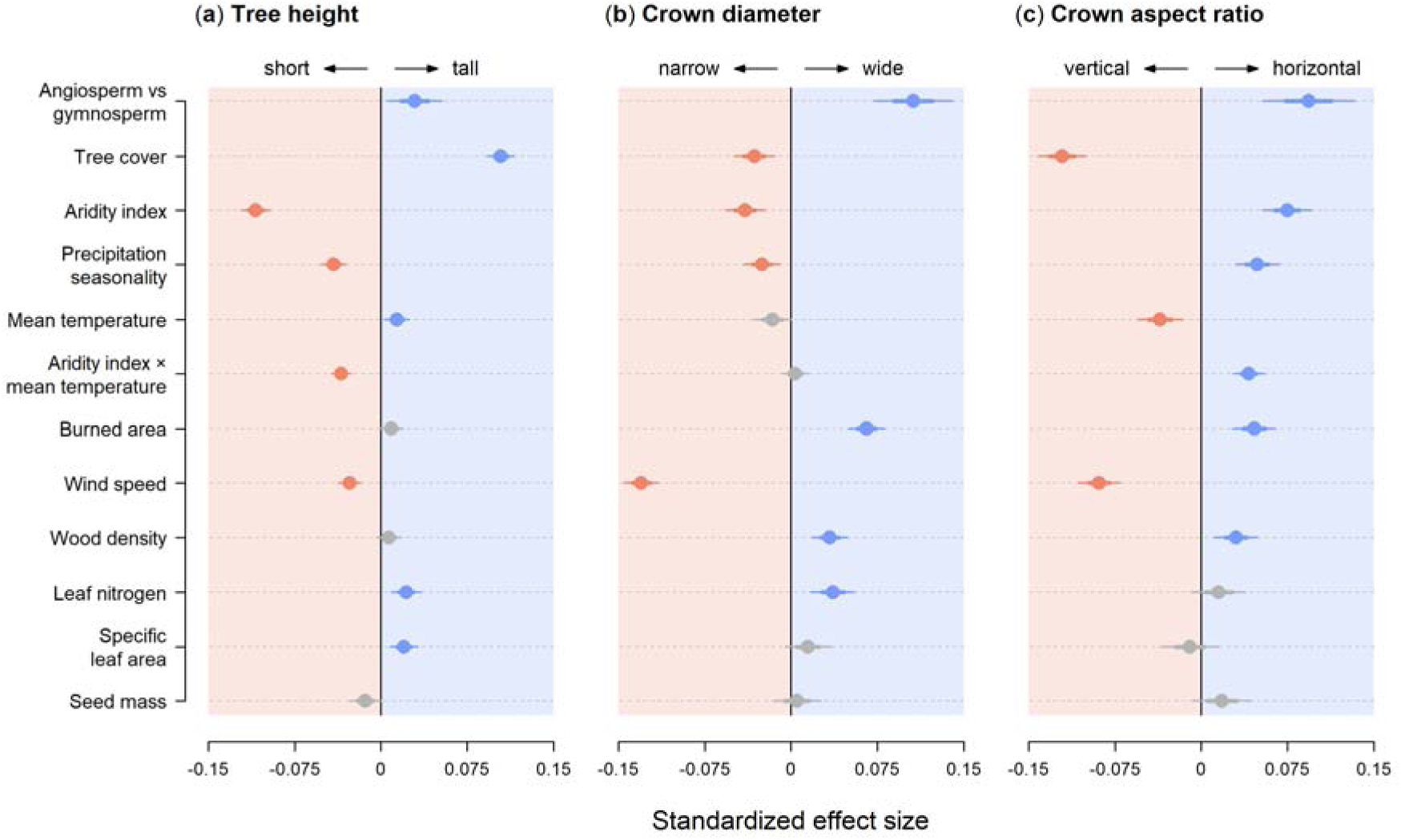
Drivers of variation in crown architecture among tree species. Points are standardized coefficients obtained from phylogenetic generalised least squares models of variation in size-standardized estimates of tree height (**a**), crown diameter (**b**) and crown aspect ratio (**c**) among species. Error bars show both standard errors (thick lines) and 95% confidence intervals (thin lines) of the model coefficients. Significantly positive and negative coefficients are shown in blue and red, respectively, while those for which the 95% confidence intervals overlap with zero are shown in grey.

**Fig. 5:**
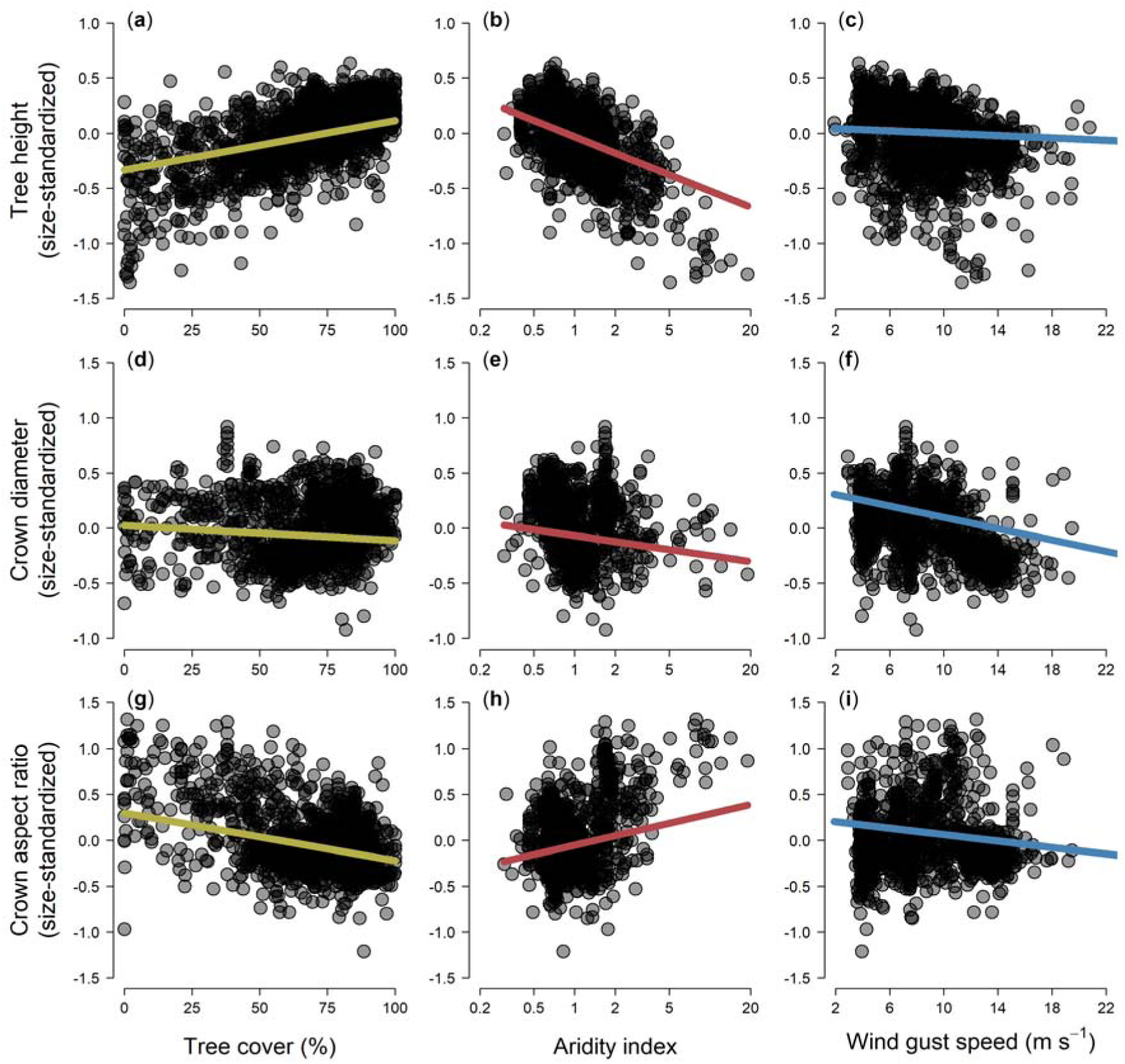
Variation in the crown architecture of tree species distributed along gradients of tree cover (left panels), aridity (middle panels) and wind gust speed (right panels). Points are species-level estimates of size-standardized tree height (*H_RESID_*, *n* = 1910 species), crown diameter (*CD_RESID_*, *n* = 1313 species) and crown aspect ratio (*CAR_RESID_*, *n* = 1309 species). Fitted lines correspond to PGLS model predictions generated by keeping all other predictors fixed at their mean values. Negative values of *H_RESID_*, *CD_RESID_*, and *CAR_RESID_*indicate species that are shorter, with narrower crowns and smaller crown aspect ratios than expected given the size of their stem, while positive values denote the opposite. Values of tree cover, aridity index and wind gust speed represent are means calculated across all individual trees of a given species. Note that the aridity index was log-transformed and that larger values correspond to drier conditions.

Water availability and tree cover also emerged as significant predictors of *CD_RESID_*, with species generally having narrower crowns in more arid and seasonal climates, and where tree cover was higher (Fig. 4b). But these effects were much less pronounced than for tree height (Fig. 5d–e). Consequently, we found that variation in *CAR_RESID_*along aridity and tree cover gradients was mostly driven by changes in *H_RESID_*, with crown profiles becoming markedly more vertical as aridity decreased and tree cover increased (Fig. 4c and Fig. 5g–h). For example, the average *CAR_D=30_* of species growing where tree cover was ≤ 20% (0.77) was more than twice that of those where tree cover was ≥ 80% (0.36). Conversely, crown diameter was much more strongly influenced by disturbances such as wind and fire (Fig. 4b). In particular, *CD_RESID_* decreased markedly as maximum wind gust speeds increased (Fig. 5f). Species also tended to be shorter for a given stem diameter where wind speeds were higher (Fig. 5c), but this effect was more subtle than for crown diameter, meaning that overall crown profiles became significantly narrower as wind speeds increased (Fig. 5i). By contrast, areas with higher frequencies of wildfires harboured species with wider crowns and higher crown aspect ratios (Fig. 4b–c), but with similar *H_RESID_* values.

While variation in *H_RESID_*, *CD_RESID_*and *CAR_RESID_* among tree species was predominantly associated with climate, tree cover and risk of disturbance, we also found that these crown architectural traits covaried with other plant functional traits (Fig. 4). After accounting for environmental effects, we found that both *H_RESID_* and *CD_RESID_* were significantly greater in species with higher leaf nitrogen content, with *H_RESID_* also positively correlated to SLA. Additionally, while we observed no clear relationships between *H_RESID_* and wood density, we did find that species with wider crowns and more horizontal crown profiles had denser wood. By contrast, none of the three crown architectural traits exhibited any relationship with seed mass.

## DISCUSSION

### Large variation in tree crown architecture across the world’s biomes and plant lineages

Using allometric data from hundreds of thousands of trees across the world, our study provides the first global picture of how tree species vary in their crown architecture (Fig. 2) and what drives this variation (Fig. 4). While ecologists have long been aware that trees can differ considerably in their crown size and shape ^1,2,5^, we have lacked a quantitative understanding of where the boundaries of this crown architectural spectrum lie. Not only did we show that tree species can vary enormously in the scaling relationships between their height, crown width and stem diameters, but we also found that size-standardised estimates of tree height (*H_RESID_*) and crown diameter (*CD_RESID_*) were largely decoupled, forming two independent axes of variation in crown architecture (Fig. 2a).

Where species fell within this crown architectural spectrum depended largely on their environment, their evolutionary history and ecological strategy. Like previous studies, we found that species confined to more arid and seasonal biomes were generally much shorter for a given stem diameter than those growing in more humid climates ^2,7,35^. We also found that the range of crown architectural types was much greater in biomes where water and temperature were non-limiting and angiosperms dominate the flora, such as tropical rainforests and temperate forests. In these environments, tree species living side by side can have incredibly different crown forms depending on their ecological strategy and evolutionary history ^7,33,50,84–86^, whereas where conditions for growth are harsher there is less flexibility in the range of crown sizes and shapes that species can assume.

From a macroevolutionary standpoint, when growing in similar environments angiosperm were only marginally taller for a given diameter than gymnosperms. However, angiosperms consistently had wider crowns. This pattern reflects a fundamental difference in the growth strategy of the two major clades, with gymnosperms investing less in lateral crown expansion due to strong apical dominance and control ^87^, resulting in crown profiles that are generally more vertical than those of angiosperms ^20,30^. When exploring how evolutionary history has shaped variation in crown architecture in more detail, we found that *H_RESID_*, *CD_RESID_* and *CAR_RESID_* all exhibited a clear phylogenetic fingerprint (Fig. 3), with several plant linages standing out. For instance, we found that dipterocarps – and trees native to Southeast Asia more generally – achieve remarkable heights for a given stem diameter ^36,37,88^. In terms of crown size, we showed that several species of Fabaceae that grow in tropical savannas in Africa and the Americas had exceptionally wide crowns. This helps explain previous observations that trees in these regions have larger crowns than those of Australia ^7,31,37^, where savannas are largely dominated by smaller-crowned eucalypts.

### Aridity and competition shape global variation in tree height scaling relationships

Aridity and tree cover emerged as the strongest predictors of tree height scaling relationships, which is consistent with previous empirical and theoretical research showing how investment in height growth is modulated by risk of hydraulic failure ^2,15,28,39,40,89^ and competition for light ^14,30,90^. Interestingly, species that were tallest for a given stem diameter (*H_D=30_* > 30 m, top 1% of species) occurred within a narrow band of aridity values (0.65–0.81) where rainfall only slightly exceeded potential evapotranspiration (Fig. 5b). Once the aridity index dipped below 0.5, species with very high *H_RESID_*values disappeared, possibly due to a combination of waterlogging from excessive rainfall and/or growth limitations linked to lower temperatures and high cloud cover ^91–93^.

*CD_RESID_* also decreased with aridity, which is consistent with trees limiting the size of their crowns (and by proxy their total leaf area) in environments where water is scarce to reduce transpiration and minimise risk of hydraulic stress ^7,30,31,94^. As expected, we also found that *CD_RESID_* decreased with tree cover, which is consistent with trees prioritizing height growth over crown expansion in response to increasing competition for light ^14,30,31^. However, the effects of both aridity and tree cover were much weaker for crown diameter than for height. One possible reason for this is that tree cover and aridity are generally negatively correlated, leading to their effects cancelling each other out: in humid climates where tree cover also tends to be high, trees would be able to support wider crowns, were it not for increased competition for space.

In contrast to aridity and tree cover, temperature played a secondary role shaping variation in tree architecture (Fig. 4). However, we did find that trees tended to be taller for a given diameter in warmer climates, which is consistent with observations that the world’s tallest trees inhabit mild and warm climates with little seasonality ^93^. This also fits our understanding of how cold temperatures impact tree height growth ^95^ and the fact that trees in cold climates generally have small vessels to minimise risk of embolisms under freezing conditions, which limits their ability to grow tall ^39^. Moreover, we found that temperature indirectly influenced tree height scaling relationships by exacerbating the effects of aridity (Fig. S5). Warmer temperatures are associated with higher vapour pressure deficits, requiring trees to have more soil water to meet higher evaporative demands ^77,96,97^. This suggests that even small decreases in rainfall and/or increases in temperature in warm climates could lead to disproportionately large impacts on forest structure ^2,98,99^.

### Disturbance regimes drive variation in crown size

Risk of disturbance by wind and fire emerged as stronger drivers of crown diameter scaling relationships than climate (Fig. 4). Species adapted to windy conditions were shorter, had narrower crowns and more vertical crown profiles for a given stem diameter than those where risk of exposure to high wind speeds was low. These adaptations would make them less prone to uprooting and snapping in high winds, as the risk of both is proportional to total crown surface area ^25,29,41,42^. Our findings are also consistent with observations that some of the world’s tallest tropical trees grow where the risk of wind disturbance is low, such as the Guiana Shield and in Borneo ^25,29,36,100^.

Crown width was also positively associated with burned area fraction, indicating that trees in fire prone environments generally allocate more resources to lateral crown expansion. This could partly be a result of lower competition for light in more open environments where fire is frequent, which would allow trees to maximise light interception by spreading their crowns laterally ^37^. It could also reflect a more direct response to fire, with trees developing wide crowns to shade out grasses and limit fuel loads ^101^. Additionally, adaptations to fire may be confounded with those associated with herbivory, which also plays an important role in shaping tree architecture in savannas ^45,46^, with wider crowns serving as a protective strategy against browsing ^37,102^. In terms of how fire might affect height scaling relationships, our expectation was that trees would generally invest more in height growth to escape fire (and herbivory). By contrast, we found no relationship between *H_RESID_* and burned area fraction, which could be because some species adopt the alternative strategy of fire resistance through the growth of thicker stems and bark ^37,45^.

As for other disturbance agents, such as snow accumulation, we would expect trees exposed to high snow loads to have narrower crowns and more slender profiles ^43,44^. While we did not test this directly, we did find that *CAR_RESID_* was generally lower in colder climates. Given that snow cover duration and mean annual temperature were highly correlated (Fig. S2), this temperature response may in part reflect an adaptation to minimising the risk of stem breakage from snow accumulation in the crown in cold climates.

### Crown architecture covaries with functional traits related to mechanical stability and photosynthesis

Crown architectural traits covaried with several other plant functional traits related to photosynthesis and structural integrity. For instance, we found that *CD_RESID_* and *CAR_RESID_* were positively associated with wood density, which is thought to confer the mechanical strength and resistance needed for trees to grow large branches and wide crowns ^8,33,34^. However, while our results support this hypothesis, we also found that numerous species had wide, horizontal crowns despite having relatively low wood densities. This highlights how other properties, such as branching architecture, may be just as important in determining a tree’s structural integrity ^25^.

Species that were taller and with wider crowns for a given stem diameter also generally had higher concentrations of nitrogen in their leaves. Leaf nitrogen content is a cornerstone of the ‘fast-slow’ plant economic spectrum ^103,104^, with species that have high leaf nitrogen generally capable of rapid growth, but also less able to tolerate shade due to higher metabolic and respiration rates ^71,72^. Based on this we would expect species with higher leaf nitrogen to invest more in both height growth and crown expansion to allow them to intercept more light, limit self-shading and optimise the distribution of leaves across their crowns – which is precisely what we found. This is also consistent with work from the U.S. showing that taller and more slender trees tend to be less shade tolerant ^35^. Similarly, early successional species in tropical forests that are adapted to grow rapidly in height to take advantage of gap openings generally have high leaf nitrogen content ^50,66^.

By contrast we found no evidence that large-seeded species were architecturally any different to those with small seeds. Seed mass has previously been shown to correlate positively with maximum plant height and canopy area, which some have proposed is the result of species with longer life spans and greater adult sizes being more likely to invest in large seeds ^70,105,106^. Alternatively, seed mass may correlate with crown architecture through its association with seed dispersal ^26,107^. For example, species with light, wind-dispersed seeds might profit from being taller to increase dispersal range, while species that have ballistic seeds or ones that simply drop to the ground might benefit from wider branches to increase distance from parent trees ^50,107^. However, we found no support for either of these hypotheses when relating variation in seed mass to *H_RESID_* and *CD_RESID_*. If these processes are at play, they may well be better captured by other facets of crown architecture (e.g., maximum tree height and crown width).

## Conclusions and future directions

Our findings highlight several fruitful avenues for future research. An obvious next step would be to expand the spectrum of crown architectural traits to other axes of crown size and shape, such as crown depth, surface area and volume ^7,14,31^. This would allow us to test long-standing predictions about how crown size and shape reflects a compromise (in terms of carbon gains) between greater light interception and higher maintenance costs ^6^, and explore how the outcome of these trade-offs varies with water and light availability ^7,108^. In this regard, efforts to better characterise crown architecture are likely to benefit from growing access to technologies such as terrestrial laser scanning (TLS). These can provide a much richer picture of a tree’s crown and local surrounding, including reconstructing its branching structure, accurate 3D volumes and within-crown distribution of leaves ^107,109,110^. Extracting these measurements from TLS point clouds remains a challenge, but access to data and automated processing pipelines are continuously improving ^1,111^.

While there is plenty of scope to build on our findings, our study provides a key starting point for characterising the crown architectural spectrum of the world’s trees. Not only do we provide the first complete picture of the range of possible crown architectural types, we also take an important step towards explaining what drives this immense variation. Our results highlight how crown architecture is jointly constrained by a range of processes related to a tree’s environment, ecological strategy and evolutionary history. This understanding will underpin ongoing efforts to leverage remote sensing technologies to track tree carbon stocks and dynamics at scale ^19–22^. It is also critical for developing the next generation of Earth system models that accurately simulate variation in vegetation structure and dynamics by incorporating more realistic representations of how tree crowns vary among biomes, plant functional types and in coordination with other traits ^15,17,18^. All of this is essential to better understanding the processes that shape the structure and function of woody biomes and tracking how these are responding to rapid global change.

## ACKNOWLEDGEMENTS

We are indebted to the countless researchers and field assistants who helped collect the field data that underpin this study and without whom this work would not have been possible. T.J. was supported by a UK NERC Independent Research Fellowship (grant: NE/S01537X/1). F.J.F. was funded by a Research Project Grant from the Leverhulme Trust (grant: RPG-2020-341). J.Ch. acknowledges an ‘Investissement d’Avenir’ grant managed by the Agence Nationale de la Recherche (CEBA grant: ANR-10-LABX-25-01 and TULIP grant: ANR-10-LABX-0041). A.A. was supported by Hebei University (grant: 521100221033), by the Jiangsu Science and Technology Special Project (grant: BX2019084), and Metasequoia Faculty Research Startup Funding at Nanjing Forestry University (grant: 163010230). G.J.L.P. was supported by projects DynAfFor (grant: CZZ1636.01D) and P3FAC (grant: CZZ1636.02D) and by the International Foundation for Science (grant: D/5822-1). T.R.F. was funded by NERC (grant: NE/N011570/1). L.F.A was supported by CAPES and ABC-CNPq (grant: 004/96). B.B.L. was supported by COMPASS-FME, a multi-institutional project supported by the U.S. Department of Energy, Office of Science, Biological and Environmental Research as part of the Environmental System Science Program. M.v.B. acknowledges funding from the Agua Salud Project, a collaboration between the Smithsonian Tropical Research Institute (STRI), the Panama Canal Authority (ACP) and the Ministry of the Environment of Panama (MiAmbiente), the Smithsonian Institution Forest Global Earth Observatory (ForestGEO), Heising-Simons Foundation, HSBC Climate Partnership, Stanley Motta, Small World Institute Fund, Frank and Kristin Levinson, the Hoch family, the U Trust, the Working Land and Seascapes Program of the Smithsonian, the National Science Foundation (grant: EAR-1360391), Singapore’s Ministry of Education and Yale–NUS College (grant: IG16-LR004). J.D. and H.L. were supported by the National Natural Science Foundation of China (grants: 41790422 and 42161144008). B.D.K. was supported by a Western Carolina University Provost Scholarship Development Grant. K.D. was supported by the African Forest Forum, and the DAAD within the framework of ClimapAfrica (Climate Research for Alumni and Postdocs in Africa) with funds of the Federal Ministry of Education and Research of Germany (grant: 91785431). H.G. was supported by a NASA GEDI Science Team Grant (grant: 80NSSC21K0201). Y.I. was supported by Grant-in-Aid for JSPS Fellows (grant: 09J04545). E.R.L. was supported by a UKRI Future Leaders Fellowship (grant: MR/T019832/1). P.M. acknowledges funding from Alex Fraser Research Forest, Forest Renewal British Columbia, and the Government of British Columbia. E.M. was supported by Swedish Energy Agency (grant: 35586-1). J.A.M. was supported by Consejo Nacional de Ciencia y Tecnología (CONACYT; grant: CB-2009-01-128136) and Universidad Nacional Autónoma de México (DGAPA-PAPIIT; grants: IN218416 and IN217620). J.M.D.R. and J.L.H.S. acknowledge funding from Reinforcing REDD+ and the South-South Cooperation Project, CONAFOR and USFS. S.C.R. acknowledges funding from FAPEMIG (grant: CAG2327-07), DAAD/CAPES and CNPq. G.S. acknowledges funding by Manchester Metropolitan University’s Environmental Science Research Centre. J.C.D. acknowledges funding from the US Department of Energy (grant: DE-SC0023309) and ANR PHYDRAUCC (grant: ANR-ANRLJ21LJCE02LJ0033LJ02). M.S. was funded by a grant from the Ministry of Education, Youth and Sports of the Czech Republic (grant: INTER-TRANSFER LTT19018). H.P. was supported by the Iran National Science Foundation (INSF). A.S. was supported by the Saxon State Ministry for Science, Culture and Tourism (grant: 3-7304/35/6-2021/48880). K.T. was supported by projects with National Institute for Environmental Studies and Hokkaido Electric Power Co., Inc, and by KAKENHI, MEXT. A.T.T. acknowledges funding from the NSF (grant: 2003205). M.A.Z. thanks the MAPA-Spain for granting access to the Spanish Forest Inventory data. The study includes data collected by partners of the official UNECE ICP Forests network (http://icp-forests.net/contributors), part of which was co-financed by the European Commission. We thank the Alberta’s Ministry of Agriculture, Forestry and Rural Economic Development for access to the Albert PSP allometric data. We are indebted to the countless researchers and field assistants who helped collect the field data compiled in the Tallo database and without whom this work would not have been possible. Dr Abd Rahman Kassim and Dr Sylvie Gourlet-Fleury, who contributed data to this project, sadly passed away before this paper was completed.

## AUTHOR CONTRIBUTIONS

TJ conceived the idea for the study. TJ led the aggregation of the data with assistance from JCh., DAC, JCa, AA, GJLP, TRF, DF and VAU, with all co-authors contributing data. TJ performed the analyses with the assistance of FJF. TJ wrote the first draft of the manuscript, with all authors providing editorial input.

## DATA AVAILABILITY STATEMENT

Data and R code supporting the results of this study will be publicly archived on Zenodo following the review of this paper. Allometry data from the TALLO database can be accessed here: https://zenodo.org/records/6637599. ICP Forests allometry data are available through request here: http://icp-forests.net/page/data-requests. Alberta PSP allometry data are available through request here: https://www.alberta.ca/permanent-sample-plots-program.aspx.

## SUPPORTING INFORMATION

Additional supporting information may be found in the online version of this article.

**Table S1**: Tree allometry database summary

**Table S2**: Climate, tree cover, disturbance and biome data sources

**Table S3**: Functional trait data sources

**Table S4**: Pairwise comparisons of *H_RESID_*, *CD_RESID_* and *CAR_RESID_* among biomes

**Table S5**: Variation in *H_RESID_*, *CD_RESID_* and *CAR_RESID_* among plant families

**Table S6**: Variation in *H_RESID_*, *CD_RESID_* and *CAR_RESID_* among plant genera

**Fig. S1**: Size-standardised estimates of tree height and crown size

**Fig. S2**: Correlations among model predictors

**Fig. S3**: MODIS-derived tree cover as a proxy for local competitive environment

**Fig. S4**: Variation in crown architectural types among gymnosperms and angiosperms

**Fig. S5**: Interactive effects of aridity and temperature on tree crown architecture

