## Supporting Information for "The global spectrum of tree crown architecture"

### Table S1 – Tree allometry database summary

**Table S1**: Number of species and trees that underpin the analyses presented in the main text. Environmental predictors included in the models were tree cover, aridity, rainfall seasonality, mean annual temperature, wind gust speed and burned area (see Table S2 for details on data sources). To test for phylogenetic signal, only species that directly matched those in the Smith & Brown (2018) phylogeny of seed plants were included.

|  | **Tree height** | | **Crown diameter** | | **Crown aspect ratio** | |
| --- | --- | --- | --- | --- | --- | --- |
| **Analysis** | **Species** | **Trees** | **Species** | **Trees** | **Species** | **Trees** |
| Environmental | 1910 | 373,666 | 1313 | 252,950 | 1309 | 251,733 |
| Environmental + wood density | 1572 | 338,925 | 1063 | 227,554 | 1059 | 226,390 |
| Environmental + leaf nitrogen | 1085 | 289,427 | 749 | 198,965 | 747 | 197,951 |
| Environmental + SLA | 1120 | 304,518 | 715 | 207,187 | 713 | 206,239 |
| Environmental + seed mass | 1108 | 311,883 | 703 | 206,347 | 701 | 205,240 |
| Phylogenetic signal | 1225 | 261,218 | 870 | 174,716 | 868 | 174,683 |

### Table S2 – Climate, tree cover, disturbance and biome data sources

**Table S2**: Sources from which data on climate, tree cover, disturbance and biome classification were obtained for this study.

| **Environmental layer** | **Units** | **Resolution** | **Format** | **Source** |
| --- | --- | --- | --- | --- |
| Mean annual temperature (MAT) | °C | 30 arc-seconds | Raster | <https://www.worldclim.org/data/worldclim21.html> |
| Maximum temperature warmest month | °C | 30 arc-seconds | Raster | <https://www.worldclim.org/data/worldclim21.html> |
| Minimum temperature coldest month | °C | 30 arc-seconds | Raster | <https://www.worldclim.org/data/worldclim21.html> |
| Temperature seasonality | °C | 30 arc-seconds | Raster | <https://www.worldclim.org/data/worldclim21.html> |
| Mean annual precipitation (MAP) | mm | 30 arc-seconds | Raster | <https://www.worldclim.org/data/worldclim21.html> |
| Precipitation seasonality | mm | 30 arc-seconds | Raster | <https://www.worldclim.org/data/worldclim21.html> |
| Potential evapotranspiration (PET) | mm | 30 arc-seconds | Raster | <https://csidotinfo.wordpress.com/2019/01/24/global-aridity-index-and-potential-evapotranspiration-climate-database-v3/> |
| Aridity index (PET/MAP) | unitless | 30 arc-seconds | Raster | <https://csidotinfo.wordpress.com/2019/01/24/global-aridity-index-and-potential-evapotranspiration-climate-database-v3/> |
| Tree cover | % | 15 arc-seconds | Raster | <https://globalmaps.github.io/ptc.html> |
| Wind gust speed ^a^ | m s^-1^ | 5 arc-minutes | Raster | <https://www.ecmwf.int/en/era5-land> |
| Burned area fraction ^b^ | % | 15 arc-minutes | Raster | <https://www.globalfiredata.org/index.html> |
| Snow cover duration ^c^ | days yr^-1^ | 15 arc-seconds | Raster | <https://download.geoservice.dlr.de/GSP/files/yearly/SCD/> |
| Terrestrial biome classification ^d^ | 7 classes† |  | Vector | <https://www.worldwildlife.org/publications/terrestrial-ecoregions-of-the-world> |

^a^ Maximum hourly wind speed between 2010-2020 estimated from ERA5-Land data

^b^ Mean burned area fraction between 2001-2010 estimated from MODIS

^c^ Maximum snow cover duration between 2001-2021 estimated from MODIS

^d^ The Terrestrial Ecoregions of the World database groups different regions of the world into 14 biomes. For the purposes of our analyses were further grouped these into 7 biome classes: **Tropical rainforests** (including a single mangrove site with analogous climate to adjacent tropical rainforests), **tropical dry forests** (combining broadleaf and coniferous forests), **temperate forests** (combining broadleaf and coniferous forests), **boreal-montane forests** (combining boreal and montane ecosystems), **tropical savannas**, **temperate woodlands** (combining Mediterranean woodlands and temperate grasslands) and **drylands**.

### Table S3 – Functional trait data sources

**Table S3**: Sources from which functional trait data were obtained for this study. Public records from the TRY database were requested on the 29/10/2020 (trait codes: 4, 14, 3117 and 26 for wood density, leaf nitrogen, SLA and seed mass, respectively). Records from the BIEN database were obtained from version 4.1.1 using the dedicated R package. Publications referenced in the table are cited in full in the main text.

| **Trait** | **Units** | **Source** | **Link** |
| --- | --- | --- | --- |
| Leaf nitrogen & SLA | mg g^-1^ & mm^2^ mg^-1^ | TRY Plant Trait Database | <https://www.try-db.org> |
| Leaf nitrogen & SLA | mg g^-1^ & mm^2^ mg^-1^ | AusTraits Database | <https://zenodo.org/records/11188867> |
| Leaf nitrogen & SLA | mg g^-1^ & mm^2^ mg^-1^ | BIEN Database | <https://bien.nceas.ucsb.edu/bien/> |
| Leaf nitrogen & SLA | mg g^-1^ & mm^2^ mg^-1^ | China Plant Trait Database | <https://esajournals.onlinelibrary.wiley.com/doi/10.1002/ecy.2091> |
| Leaf nitrogen & SLA | mg g^-1^ & mm^2^ mg^-1^ | Terrestrial Ecosystem Research Network | <https://supersites.tern.org.au/knb/metacat/supersite.949.4/html> |
| Leaf nitrogen & SLA | mg g^-1^ & mm^2^ mg^-1^ | Both *et al*. (2019) | <https://zenodo.org/records/3247631> |
| Leaf nitrogen & SLA | mg g^-1^ & mm^2^ mg^-1^ | Unpublished data from Iran |  |
| Seed mass | g | Kew Gardens Seed Information Database | <http://data.kew.org/sid/> |
| Seed mass | g | TRY Plant Trait Database | <https://www.try-db.org> |
| Seed mass | g | AusTraits Database | <https://doi.org/10.5281/zenodo.3568429> |
| Seed mass | g | Unpublished data from Hainan |  |
| Seed mass | g | Unpublished data from Iran |  |
| Wood density | g cm^-3^ | Global Wood Density Database | <https://datadryad.org/stash/dataset/doi:10.5061/dryad.234> |
| Wood density | g cm^-3^ | TRY Plant Trait Database | <https://www.try-db.org> |
| Wood density | g cm^-3^ | AusTraits Database | <https://zenodo.org/records/11188867> |
| Wood density | g cm^-3^ | Brown *et al*. (1997) | <https://www.fao.org/4/w4095e/w4095e0c.htm> |
| Wood density | g cm^-3^ | Díaz *et al*. (2015) | <http://www.scielo.org.mx/pdf/mb/v21nspe/v21nspea6.pdf> |
| Wood density | g cm^-3^ | Bradford *et al*. (2014) | <https://supersites.tern.org.au/knb/metacat/supersite.174/html> |
| Wood density | g cm^-3^ | Mori *et al*. (2014) | <https://academic.oup.com/jpe/article/7/4/356/977001> |
| Wood density | g cm^-3^ | Iida *et al*. (2012) | <https://doi.org/10.1111/j.1365-2435.2011.01921.x> |
| Wood density | g cm^-3^ | Unpublished data from Iran |  |

### Table S4 – Pairwise comparisons of *H_RESID_*, *CD_RESID_* and *CAR_RESID_* among biomes

**Table S4**: Pairwise differences in size-standardized estimates of tree height (*H_RESID_*), crown diameter (*CD_RESID_*) and crown aspect ratio (*CAR_RESID_*) among biomes. Differences among biomes were tested using one-way ANOVAs with *post hoc* Tukey tests. Biome association explained 33%, 5% and 39% of the variation in *H_RESID_*, *CD_RESID_* and *CAR_RESID_* among species, respectively. Statistically significant differences among biomes are highlighted in bold (*P* < 0.05).

| **Pairwise comparison** | **Difference in *H_RESID_*** | ***P*-value** | **Difference in *CD_RESID_*** | ***P*-value** | **Difference in *CAR_RESID_*** | ***P*-value** |
| --- | --- | --- | --- | --- | --- | --- |
| Dryland vs Boreal-montane forest | **-1.40** | **<0.0001** | -0.02 | 1.000 | 1.44 | **<0.0001** |
| Temperate forest vs Boreal-montane forest | -0.15 | 0.522 | 0.13 | 0.824 | 0.30 | **0.026** |
| Temperate forest vs Dryland | **1.24** | **<0.0001** | 0.15 | 0.949 | -1.14 | **<0.0001** |
| Temperate woodland vs Boreal-montane forest | **-0.51** | **<0.0001** | 0.16 | 0.720 | 0.74 | **<0.0001** |
| Temperate woodland vs Dryland | **0.88** | **<0.0001** | 0.18 | 0.897 | -0.70 | **<0.0001** |
| Temperate woodland vs Temperate forest | **-0.36** | **<0.0001** | 0.03 | 0.996 | 0.44 | **<0.0001** |
| Tropical dry forest vs Boreal-montane forest | -0.24 | 0.081 | **0.37** | **0.006** | 0.68 | **<0.0001** |
| Tropical dry forest vs Dryland | **1.16** | **<0.0001** | 0.38 | 0.121 | -0.76 | **<0.0001** |
| Tropical dry forest vs Temperate forest | -0.09 | 0.193 | **0.24** | **<0.0001** | 0.37 | **<0.0001** |
| Tropical dry forest vs Temperate woodland | **0.27** | **<0.0001** | **0.21** | **0.016** | -0.06 | 0.946 |
| Tropical rainforest vs Boreal-montane forest | -0.04 | 0.999 | 0.11 | 0.911 | 0.20 | 0.319 |
| Tropical rainforest vs Dryland | **1.35** | **<0.0001** | 0.12 | 0.976 | -1.24 | **<0.0001** |
| Tropical rainforest vs Temperate forest | **0.11** | **<0.0001** | -0.02 | 0.961 | -0.10 | **<0.0001** |
| Tropical rainforest vs Temperate woodland | **0.47** | **<0.0001** | -0.05 | 0.929 | -0.54 | **<0.0001** |
| Tropical rainforest vs Tropical dry forest | **0.20** | **<0.0001** | **-0.26** | **<0.0001** | -0.48 | **<0.0001** |
| Tropical savanna vs Boreal-montane forest | **-0.58** | **<0.0001** | 0.34 | **0.013** | 0.97 | **<0.0001** |
| Tropical savanna vs Dryland | **0.82** | **<0.0001** | 0.35 | 0.187 | -0.47 | **0.019** |
| Tropical savanna vs Temperate forest | **-0.43** | **<0.0001** | **0.21** | **<0.0001** | 0.67 | **<0.0001** |
| Tropical savanna vs Temperate woodland | -0.07 | 0.817 | **0.17** | **0.043** | 0.23 | **0.002** |
| Tropical savanna vs Tropical dry forest | **-0.34** | **<0.0001** | -0.03 | 0.995 | 0.29 | **<0.0001** |
| Tropical savanna vs Tropical rainforest | **-0.54** | **<0.0001** | **0.23** | **<0.0001** | 0.77 | **<0.0001** |

### Table S5 – Variation in *H_RESID_*, *CD_RESID_* and *CAR_RESID_* among plant families

**Table S5**: Mean values of size-standardized estimates of tree height (*H_RESID_*), crown diameter (*CD_RESID_*) and crown aspect ratio (*CAR_RESID_*) of plant families represented by at least 5 species in the analysis (*n* = 63 families for *H****_RESID_*** and 56 families for *CD****_RESID_*** and *CAR****_RESID_***). An ANOVA fit without an intercept was used to test whether family-level mean values were significantly different from zero (*P* < 0.05, highlighted in bold).

| **Family** | ***H_RESID_*** | ***P*-value** | ***CD_RESID_*** | ***P*-value** | ***CAR_RESID_*** | ***P*-value** |
| --- | --- | --- | --- | --- | --- | --- |
| Achariaceae | 0.043 | 0.652 |  |  |  |  |
| Altingiaceae | -0.026 | 0.828 | -0.104 | 0.392 | -0.082 | 0.580 |
| Anacardiaceae | **-0.086** | **0.048** | -0.018 | 0.735 | 0.114 | 0.087 |
| Annonaceae | **0.176** | **0.000** | **0.112** | **0.022** | -0.064 | 0.286 |
| Apocynaceae | 0.051 | 0.312 | -0.049 | 0.396 | -0.091 | 0.202 |
| Aquifoliaceae | -0.040 | 0.589 | -0.069 | 0.400 | -0.037 | 0.711 |
| Araliaceae | 0.017 | 0.831 | 0.013 | 0.892 | 0.064 | 0.588 |
| Betulaceae | **0.159** | **0.005** | **0.159** | **0.013** | -0.011 | 0.885 |
| Bignoniaceae | -0.008 | 0.920 | -0.127 | 0.185 | -0.137 | 0.244 |
| Boraginaceae | -0.002 | 0.987 | 0.018 | 0.862 | 0.030 | 0.814 |
| Burseraceae | **0.188** | **0.000** | 0.053 | 0.352 | **-0.179** | **0.010** |
| Calophyllaceae | **0.154** | **0.038** | **0.223** | **0.014** | -0.019 | 0.864 |
| Cannabaceae | 0.018 | 0.798 | 0.038 | 0.610 | 0.006 | 0.946 |
| Celastraceae | 0.039 | 0.647 | 0.025 | 0.821 | -0.081 | 0.550 |
| Chrysobalanaceae | **0.242** | **0.011** | 0.214 | 0.054 | -0.013 | 0.923 |
| Clusiaceae | 0.066 | 0.395 | 0.151 | 0.095 | 0.081 | 0.465 |
| Combretaceae | **-0.298** | **0.000** | **0.165** | **0.010** | **0.495** | **0.000** |
| Cornaceae | 0.093 | 0.396 | 0.050 | 0.680 | -0.004 | 0.979 |
| Cunoniaceae | 0.114 | 0.201 |  |  |  |  |
| Cupressaceae | **-0.277** | **0.000** | **-0.188** | **0.022** | 0.104 | 0.302 |
| Dipterocarpaceae | **0.262** | **0.000** | 0.056 | 0.220 | **-0.236** | **0.000** |
| Ebenaceae | -0.001 | 0.987 | -0.105 | 0.120 | -0.098 | 0.238 |
| Elaeocarpaceae | 0.087 | 0.114 | **-0.187** | **0.030** | -0.086 | 0.412 |
| Ericaceae | **-0.283** | **0.003** | 0.057 | 0.604 | 0.262 | 0.055 |
| Euphorbiaceae | **0.080** | **0.018** | 0.022 | 0.594 | -0.078 | 0.117 |
| Fabaceae | **-0.085** | **0.000** | **0.168** | **0.000** | **0.241** | **0.000** |
| Fagaceae | **-0.124** | **0.000** | -0.019 | 0.500 | **0.099** | **0.005** |
| Juglandaceae | 0.003 | 0.971 | 0.133 | 0.196 | 0.106 | 0.399 |
| Lamiaceae | -0.157 | 0.064 | **-0.284** | **0.020** | -0.264 | 0.077 |
| Lauraceae | **0.085** | **0.001** | **-0.161** | **0.000** | **-0.159** | **0.000** |
| Lecythidaceae | **0.150** | **0.021** | 0.025 | 0.747 | **-0.260** | **0.007** |
| Loganiaceae | **-0.459** | **0.000** | 0.202 | 0.095 | **0.667** | **0.000** |
| Magnoliaceae | -0.042 | 0.622 | **-0.232** | **0.024** | -0.156 | 0.216 |
| Malvaceae | -0.005 | 0.879 | -0.043 | 0.224 | -0.045 | 0.302 |
| Melastomataceae | 0.088 | 0.386 | 0.133 | 0.229 | -0.011 | 0.936 |
| Meliaceae | **0.083** | **0.041** | 0.031 | 0.499 | -0.041 | 0.474 |
| Moraceae | -0.038 | 0.381 | -0.081 | 0.129 | -0.013 | 0.841 |
| Myristicaceae | **0.253** | **0.000** | **0.148** | **0.025** | -0.130 | 0.108 |
| Myrtaceae | **-0.054** | **0.005** | **-0.106** | **0.004** | -0.087 | 0.057 |
| Nothofagaceae | -0.070 | 0.562 |  |  |  |  |
| Nyctaginaceae | -0.155 | 0.196 |  |  |  |  |
| Olacaceae | **0.167** | **0.039** | **0.182** | **0.034** | 0.014 | 0.895 |
| Oleaceae | -0.016 | 0.852 | -0.081 | 0.343 | -0.084 | 0.423 |
| Pentaphylacaceae | -0.035 | 0.683 | -0.150 | 0.097 | -0.102 | 0.359 |
| Phyllanthaceae | -0.031 | 0.528 | 0.008 | 0.881 | 0.049 | 0.479 |
| Pinaceae | **-0.121** | **0.000** | **-0.185** | **0.000** | **-0.136** | **0.002** |
| Podocarpaceae | -0.072 | 0.418 | -0.173 | 0.071 | -0.101 | 0.390 |
| Primulaceae | -0.071 | 0.426 | **-0.267** | **0.005** | -0.196 | 0.097 |
| Proteaceae | 0.085 | 0.157 |  |  |  |  |
| Putranjivaceae | **0.189** | **0.020** | **0.213** | **0.013** | -0.033 | 0.754 |
| Rhamnaceae | 0.054 | 0.567 | 0.060 | 0.621 | 0.180 | 0.227 |
| Rosaceae | -0.110 | 0.091 | -0.047 | 0.515 | 0.117 | 0.188 |
| Rubiaceae | -0.004 | 0.931 | **-0.116** | **0.027** | **-0.136** | **0.034** |
| Rutaceae | 0.069 | 0.133 | -0.086 | 0.256 | -0.048 | 0.605 |
| Salicaceae | **0.112** | **0.040** | 0.018 | 0.779 | -0.082 | 0.298 |
| Sapindaceae | 0.052 | 0.172 | -0.012 | 0.795 | -0.033 | 0.574 |
| Sapotaceae | **0.118** | **0.007** | 0.058 | 0.284 | -0.110 | 0.099 |
| Symplocaceae | -0.096 | 0.216 | -0.106 | 0.176 | -0.001 | 0.995 |
| Theaceae | -0.064 | 0.447 | **-0.211** | **0.014** | -0.144 | 0.171 |
| Thymelaeaceae | 0.105 | 0.381 |  |  |  |  |
| Ulmaceae | -0.120 | 0.156 | **0.268** | **0.003** | **0.393** | **0.000** |
| Urticaceae | **0.211** | **0.009** | 0.115 | 0.233 | -0.126 | 0.286 |
| Vochysiaceae | 0.053 | 0.659 |  |  |  |  |

### Table S6 – Variation in *H_RESID_*, *CD_RESID_* and *CAR_RESID_* among plant genera

**Table S6**: Mean values of size-standardized estimates of tree height (*H_RESID_*), crown diameter (*CD_RESID_*) and crown aspect ratio (*CAR_RESID_*) of plant genera represented by at least 5 species in the analysis (*n* = 85 genera for *H_resid_* and 60 genera for *CD_resid_* and *CAR_resid_*). An ANOVA fit without an intercept was used to test whether genus-level mean values were significantly different from zero (*P* < 0.05, highlighted in bold).

| **Genus** | ***H_resid_*** | ***P*-value** | ***CD_resid_*** | ***P*-value** | ***CAR_resid_*** | ***P*-value** |
| --- | --- | --- | --- | --- | --- | --- |
| Abies | **-0.169** | **0.016** | **-0.243** | **0.004** | **-0.238** | **0.016** |
| Acacia | **-0.303** | **0.000** | **0.397** | **0.000** | **0.715** | **0.000** |
| Acer | 0.043 | 0.478 | 0.057 | 0.394 | -0.003 | 0.971 |
| Aglaia | 0.106 | 0.330 | 0.002 | 0.985 | -0.106 | 0.420 |
| Alangium | 0.119 | 0.273 |  |  |  |  |
| Albizia | **-0.231** | **0.004** | 0.077 | 0.355 | **0.314** | **0.001** |
| Alnus | 0.162 | 0.137 |  |  |  |  |
| Alstonia | 0.102 | 0.346 |  |  |  |  |
| Archidendron | -0.027 | 0.800 |  |  |  |  |
| Artocarpus | 0.041 | 0.636 | -0.123 | 0.225 | -0.160 | 0.183 |
| Aspidosperma | **0.262** | **0.016** | -0.004 | 0.975 | -0.255 | 0.053 |
| Beilschmiedia | **0.141** | **0.030** | -0.132 | 0.133 | -0.191 | 0.067 |
| Betula | 0.143 | 0.062 | 0.126 | 0.181 | -0.036 | 0.748 |
| Brachystegia | **-0.310** | **0.004** | **0.317** | **0.005** | **0.630** | **0.000** |
| Calophyllum | 0.144 | 0.075 | **0.209** | **0.027** | -0.016 | 0.884 |
| Canarium | 0.145 | 0.114 | -0.121 | 0.275 | **-0.264** | **0.046** |
| Carya | 0.137 | 0.206 |  |  |  |  |
| Castanopsis | -0.109 | 0.083 | **-0.152** | **0.028** | -0.005 | 0.950 |
| Cecropia | 0.209 | 0.054 |  |  |  |  |
| Celtis | 0.075 | 0.304 | 0.096 | 0.249 | -0.016 | 0.868 |
| Chrysophyllum | 0.124 | 0.209 | **0.305** | **0.006** | 0.172 | 0.192 |
| Cinnamomum | -0.010 | 0.906 | **-0.256** | **0.012** | -0.211 | 0.080 |
| Combretum | **-0.512** | **0.000** | **0.203** | **0.046** | **0.795** | **0.000** |
| Cordia | 0.051 | 0.605 | 0.042 | 0.704 | -0.005 | 0.969 |
| Corymbia | -0.089 | 0.108 |  |  |  |  |
| Croton | 0.130 | 0.130 | 0.164 | 0.108 | 0.006 | 0.958 |
| Cryptocarya | **0.118** | **0.029** | **-0.184** | **0.027** | -0.123 | 0.211 |
| Diospyros | -0.001 | 0.985 | -0.105 | 0.090 | -0.098 | 0.182 |
| Dipterocarpus | **0.281** | **0.005** |  |  |  |  |
| Drypetes | **0.189** | **0.010** | **0.213** | **0.007** | -0.033 | 0.723 |
| Dysoxylum | 0.089 | 0.367 |  |  |  |  |
| Elaeocarpus | 0.069 | 0.242 | **-0.299** | **0.002** | -0.153 | 0.170 |
| Endiandra | **0.300** | **0.000** |  |  |  |  |
| Eucalyptus | **-0.116** | **0.000** | **-0.207** | **0.006** | -0.008 | 0.929 |
| Fagus | 0.088 | 0.417 | **0.301** | **0.007** | 0.154 | 0.242 |
| Ficus | **-0.165** | **0.019** | **-0.264** | **0.002** | -0.020 | 0.842 |
| Flindersia | **0.346** | **0.000** |  |  |  |  |
| Fraxinus | 0.015 | 0.880 | 0.083 | 0.414 | 0.030 | 0.805 |
| Garcinia | 0.086 | 0.288 | 0.184 | 0.070 | 0.085 | 0.482 |
| Glochidion | 0.100 | 0.314 |  |  |  |  |
| Heritiera | 0.119 | 0.273 |  |  |  |  |
| Hopea | 0.192 | 0.053 | -0.110 | 0.325 | **-0.269** | **0.042** |
| Hydnocarpus | -0.004 | 0.968 |  |  |  |  |
| Ilex | -0.040 | 0.551 | -0.069 | 0.359 | -0.037 | 0.676 |
| Inga | **0.250** | **0.022** | **0.259** | **0.020** | 0.007 | 0.959 |
| Knema | **0.291** | **0.002** | **0.248** | **0.026** | -0.068 | 0.605 |
| Larix | 0.133 | 0.181 | -0.132 | 0.236 | -0.250 | 0.059 |
| Licania | **0.282** | **0.009** |  |  |  |  |
| Lindera | -0.046 | 0.641 | -0.185 | 0.070 | -0.128 | 0.288 |
| Lithocarpus | -0.011 | 0.846 | -0.103 | 0.090 | -0.087 | 0.221 |
| Litsea | 0.111 | 0.098 | -0.057 | 0.472 | -0.076 | 0.417 |
| Lonchocarpus | -0.006 | 0.947 | 0.184 | 0.099 | 0.212 | 0.109 |
| Macaranga | 0.103 | 0.140 | -0.035 | 0.710 | -0.155 | 0.165 |
| Machilus | -0.104 | 0.199 | **-0.291** | **0.000** | -0.177 | 0.071 |
| Magnolia | -0.079 | 0.330 | **-0.301** | **0.003** | -0.159 | 0.188 |
| Mallotus | -0.008 | 0.938 |  |  |  |  |
| Manilkara | 0.052 | 0.634 |  |  |  |  |
| Melaleuca | **-0.427** | **0.000** |  |  |  |  |
| Myristica | **0.229** | **0.021** |  |  |  |  |
| Neolitsea | -0.033 | 0.697 | **-0.300** | **0.003** | **-0.239** | **0.048** |
| Nothofagus | -0.070 | 0.522 |  |  |  |  |
| Ocotea | **0.203** | **0.041** |  |  |  |  |
| Ormosia | 0.040 | 0.665 | **-0.206** | **0.043** | -0.215 | 0.074 |
| Palaquium | **0.262** | **0.016** |  |  |  |  |
| Picea | 0.054 | 0.484 | **-0.299** | **0.001** | **-0.412** | **0.000** |
| Pinus | **-0.208** | **0.000** | **-0.142** | **0.002** | 0.007 | 0.893 |
| Populus | 0.131 | 0.154 | 0.061 | 0.519 | -0.073 | 0.514 |
| Pouteria | **0.246** | **0.007** | 0.045 | 0.634 | -0.217 | 0.052 |
| Protium | **0.280** | **0.002** | 0.160 | 0.152 | -0.116 | 0.377 |
| Prunus | 0.038 | 0.698 | 0.056 | 0.615 | 0.075 | 0.570 |
| Pterocarpus | **-0.200** | **0.030** | **0.297** | **0.004** | **0.506** | **0.000** |
| Quercus | **-0.178** | **0.000** | 0.007 | 0.838 | **0.178** | **0.000** |
| Schima | -0.020 | 0.851 | -0.108 | 0.331 | -0.091 | 0.491 |
| Shorea | **0.282** | **0.000** | 0.009 | 0.894 | **-0.348** | **0.000** |
| Sloanea | 0.129 | 0.194 |  |  |  |  |
| Sterculia | 0.182 | 0.095 | -0.015 | 0.896 | -0.200 | 0.129 |
| Strychnos | **-0.459** | **0.000** | 0.202 | 0.069 | **0.667** | **0.000** |
| Symplocos | -0.096 | 0.173 | -0.106 | 0.140 | -0.001 | 0.994 |
| Syzygium | 0.061 | 0.083 | -0.071 | 0.122 | -0.102 | 0.063 |
| Terminalia | **-0.188** | **0.010** | 0.147 | 0.118 | **0.289** | **0.010** |
| Trichilia | 0.157 | 0.148 | **0.360** | **0.001** | 0.213 | 0.107 |
| Ulmus | -0.166 | 0.093 | **0.219** | **0.031** | **0.408** | **0.001** |
| Vachellia | **-0.768** | **0.000** | 0.061 | 0.514 | **0.975** | **0.000** |
| Virola | **0.272** | **0.006** |  |  |  |  |
| Vitex | -0.207 | 0.057 |  |  |  |  |

### Fig. S1 – Size-standardised estimates of tree height and crown size

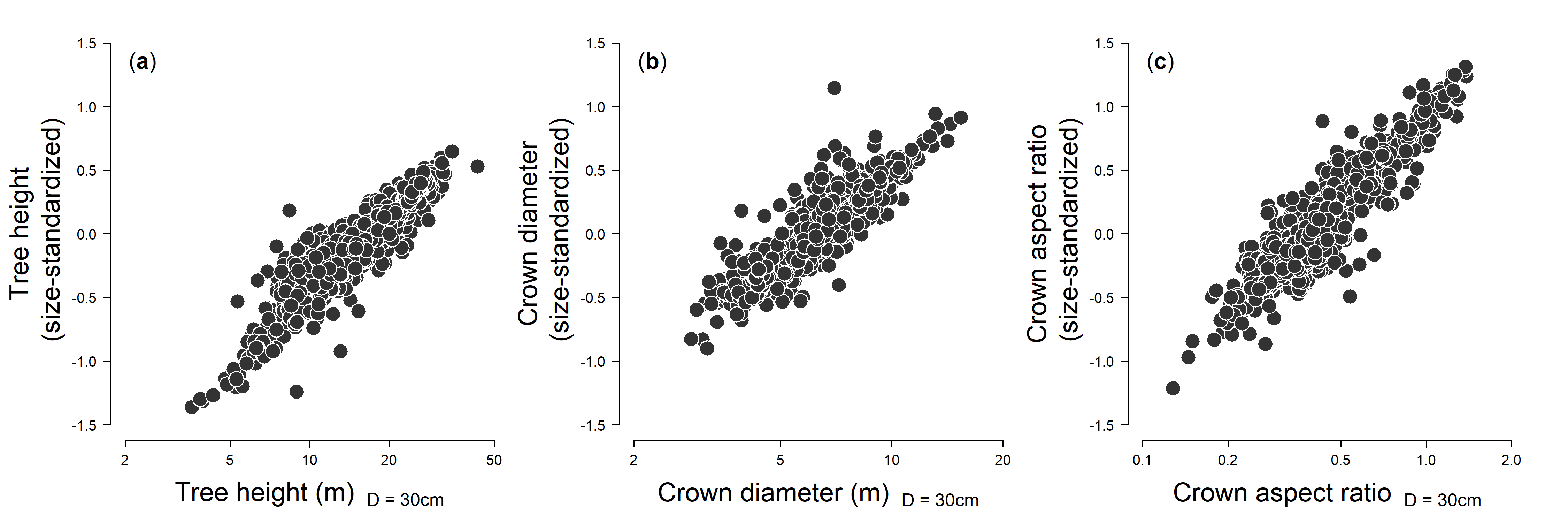

**Fig. S1**: Comparison between species’ size-standardized estimates of (**a**) tree height, (**b**) crown diameter and (**c**) crown aspect ratio generated using the model residual approach described in the main text and predicted values of all three attributes estimated for a tree of fixed size (stem diameter, *D* = 30 cm). Pearson’s correlation coefficients for the comparisons shown above ranged between 0.91–0.93 (*P* < 0.0001 in all cases).

### Fig. S2 – Correlations among model predictors

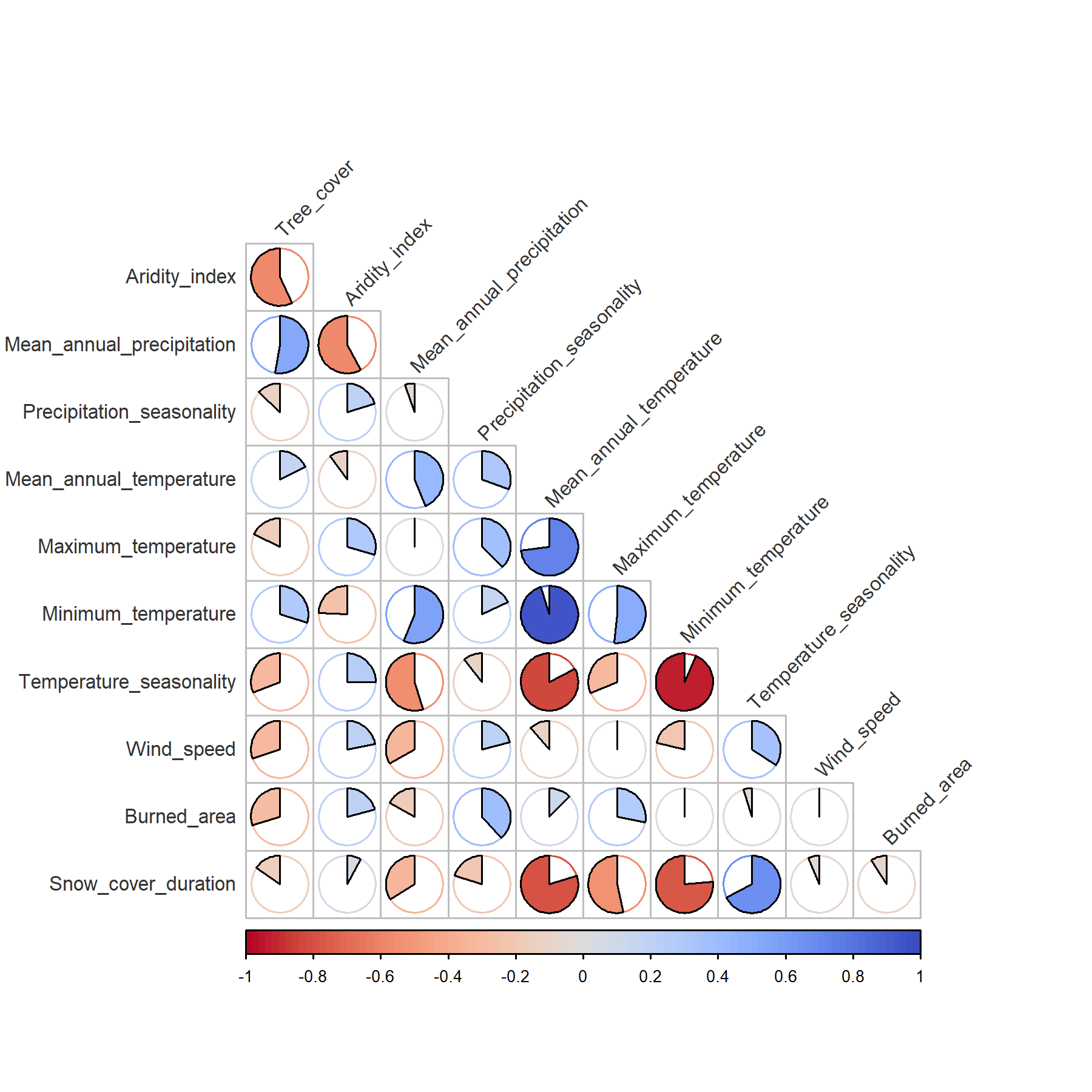

**Fig. S2**: Pearson’s correlation coefficients among bioclimatic and disturbance predictors used in the models described in the main text. Data sources for each bioclimatic and disturbance attribute are reported in Table S2.

### Fig. S3 – MODIS-derived tree cover as a proxy for local competitive environment

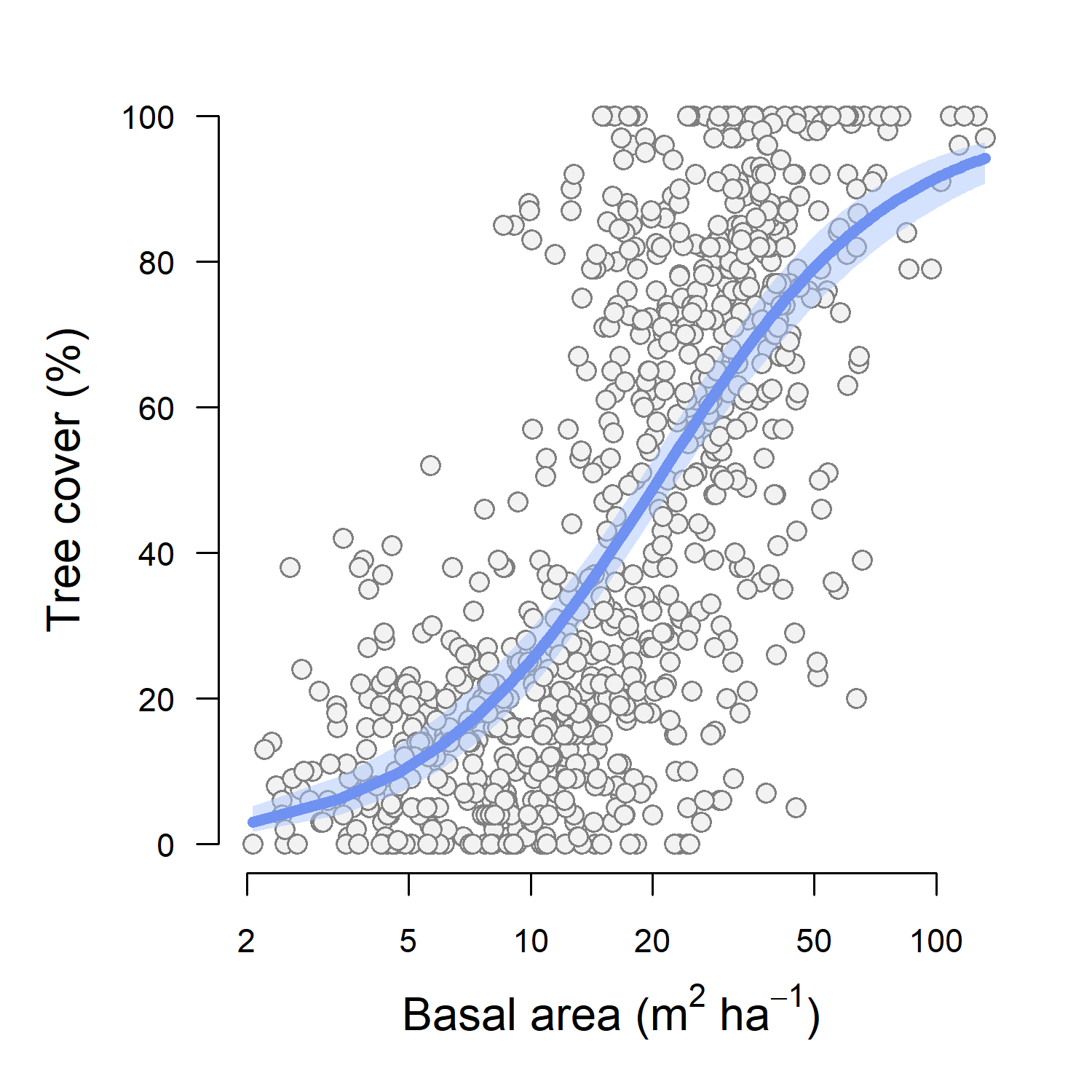

**Fig. S3**: Relationship between MODIS-derived estimates of tree cover and field-measured values of basal area across 851 forest plots. A line of best fit with shaded 95% confidence intervals from a binomial GLM is shown in blue. We used data from 851 geo-located forest plots spanning major forested and non-forested biomes to evaluate how well MODIS-derived estimates of tree cover compare to field-estimated proxies of competitive environment – specifically forest basal area. Tree cover was estimated from MODIS at 500 m resolution for the year 2008 (Kobayashi *et al.*, 2016). Plot-level basal area values were obtained from global and regional databases, including the Forest Observation System ([www.forest-observation-system.net](http://www.forest-observation-system.net)) (Schepaschenko *et al.*, 2019), the FunDivEUROPE network (Baeten *et al.*, 2013), Australia’s Biomass Plot Library ([www.auscover.org.au/datasets/biomass-plot-library](http://www.auscover.org.au/datasets/biomass-plot-library)), and a network of forest plots distributed across the Gola Rainforest National Park in Sierra Leone (Jucker *et al.*, 2016). Using these data, we fit a binomial GLM relating variation in tree cover (scaled between 0–1) to plot-level estimates of basal area (log-transformed). This revealed a significantly positive relationship between tree cover and forest basal area (*P* < 0.0001, McFadden’s pseudo *R^2^* = 0.48; Supplementary Fig. 3), suggesting that MODIS-derived estimates of tree cover provide a robust proxy of a tree’s local competitive environment.

### Fig. S4 – Variation in crown architectural types among gymnosperms and angiosperms

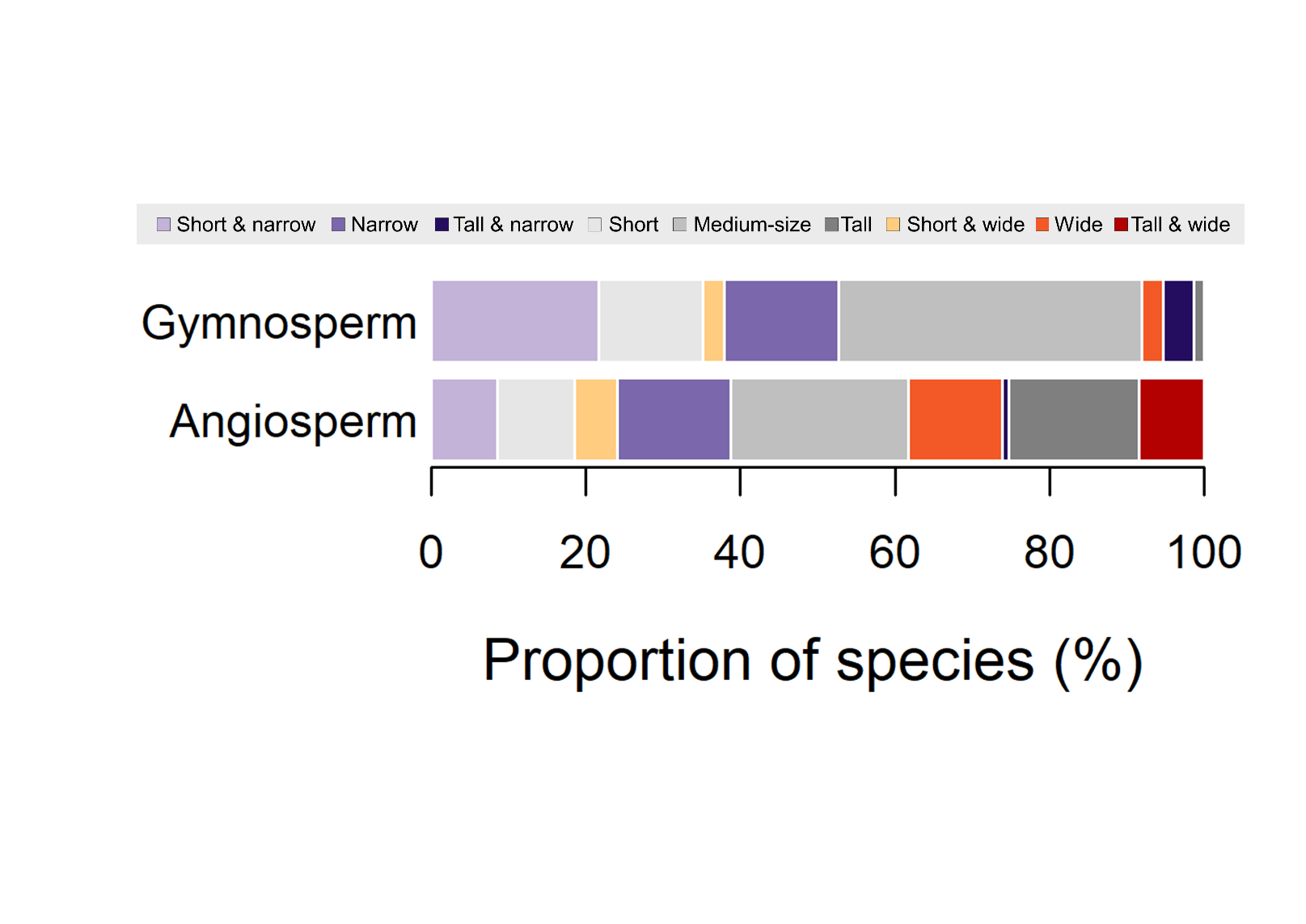

**Fig. S4**: Distribution of tree crown architectural types among gymnosperms and angiosperms for the 1309 tree species for which both height and crown size were measured. Tree species were grouped into one of nine architectural types based on their size-standardized height and crown diameter values (see Fig. 2 in the main text for details).

### Fig. S5 – Interactive effects of aridity and temperature on tree crown architecture

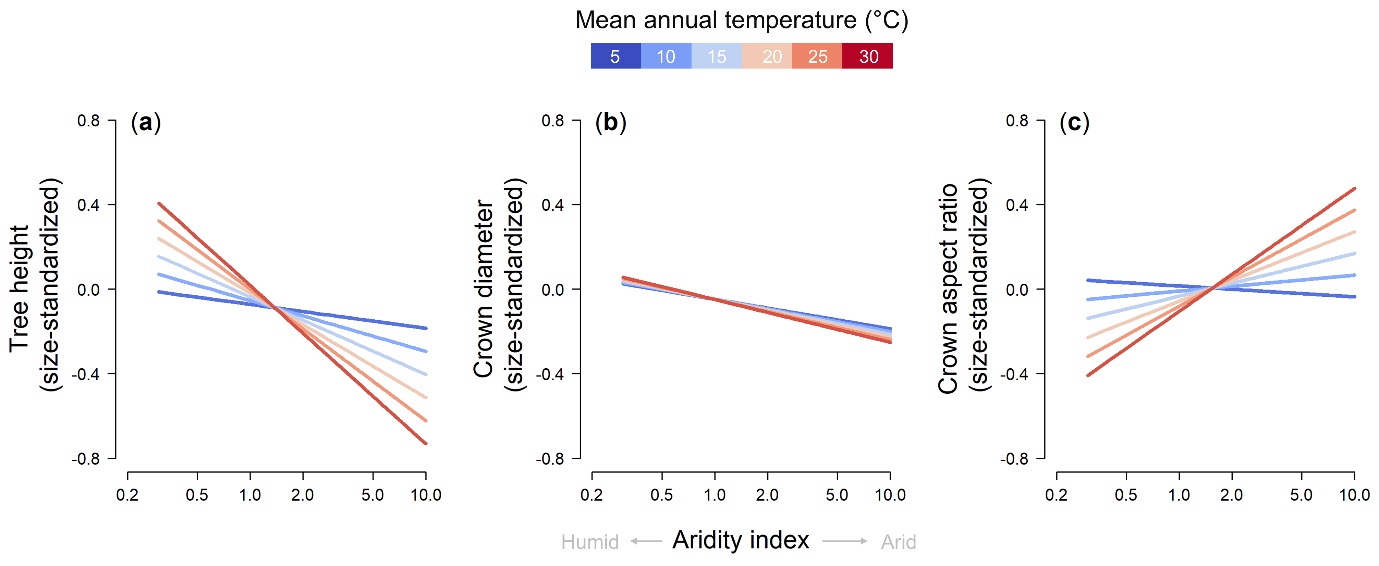

**Fig. S5**: Interactive effects of aridity and mean annual temperature on tree crown architecture. Lines show how species’ size-standardized height (**a**), crown diameter (**b**) and crown aspect ratio (**c**) values are predicted to vary along an aridity gradient for different levels of mean annual temperature. Fitted lines were generated from the phylogenetic generalised least squares models while keeping all other predictors fixed at their mean value.
